# Lineage plasticity in SCLC generates non-neuroendocrine cells primed for vasculogenic mimicry

**DOI:** 10.1101/2022.10.21.512986

**Authors:** Sarah M Pearsall, Stuart C Williamson, Fernando J García Marqués, Sam Humphrey, Ellyn Hughes, Yan Ting Shue, Abel Bermudez, Kristopher K Frese, Melanie Galvin, Mathew Carter, Lynsey Priest, Alastair Kerr, Cong Zhou, Trudy G. Oliver, Jonathan D Humphries, Martin J. Humphries, Fiona Blackhall, Ian G Cannell, Sharon J Pitteri, Gregory J Hannon, Julien Sage, Kathryn L Simpson, Caroline Dive

## Abstract

**Introduction:** Vasculogenic mimicry (VM), the process of tumor cell trans-differentiation to endow endothelial-like characteristics supporting *de novo* vessel formation, is associated with poor prognosis in several tumor types, including small cell lung cancer (SCLC). In genetically engineered mouse models (GEMMs) of SCLC, NOTCH and MYC co-operate to drive a neuroendocrine (NE) to non-NE phenotypic switch and co-operation between NE and non-NE cells is required for metastasis. Here, we define the phenotype of VM-competent cells and molecular mechanisms underpinning SCLC VM using circulating tumor cell-derived explant (CDX) models and GEMMs.

**Methods:** We analysed perfusion within VM vessels and their association with NE and non-NE phenotypes using multiplex immunohistochemistry in CDX and GEMMs. VM-proficient cell subpopulations in *ex vivo* cultures were molecularly profiled by RNA sequencing and mass spectrometry. We evaluated their 3D structure and defined collagen-integrin interactions.

**Results:** We show that VM vessels are present in 23/25 CDX models and in 2 GEMMs. Perfused VM vessels support tumor growth and only Notch-active non-NE cells are VM-competent *in vivo* and *ex vivo*, expressing pseudohypoxia, blood vessel development and extracellular matrix (ECM) organization signatures. On Matrigel, VM-primed non-NE cells re-model ECM into hollow tubules in an integrin β1-dependent process.

**Conclusions:** We identify VM as an exemplar of functional heterogeneity and plasticity in SCLC and these findings take significant steps towards understanding the molecular events that enable VM. These results support therapeutic co-targeting of both NE and non-NE cells to curtail SCLC progression and to improve SCLC patient outcomes in future.

## Introduction

SCLC patients typically present with high circulating tumor cell (CTC) burden and early, widespread metastasis with a 5-year survival of <7%^1,2^. Despite inter and intra-tumor heterogeneity, SCLC treatment is homogeneous (platinum-etoposide chemotherapy) and responses are short-lived^2^. Immunotherapy was recently incorporated into standard of care; albeit benefiting only ∼15% people within an unselected subpopulation^3-5^. As research biopsies present a significant challenge, we pioneered the generation of CTC-derived eXplants (CDX) from peripheral blood^6^. CDX faithfully recapitulate the histopathology, recently defined molecular subtypes^7^ and chemotherapy responses of donor patient tumors^8^. In SCLC GEMMs, NOTCH and MYC co-operate to drive phenotype switching from NE to non-NE cells^9-11^ where non-NE cells are less tumorigenic but support NE cell expansion *in vivo*^12^, and where paracrine signaling between NE and non-NE cells facilitates metastasis^13^. Functional plasticity accompanied by increased intra-tumoral heterogeneity and epithelial to mesenchymal transition with loss of NE phenotype is observed during chemotherapy resistance^14^. Induction of the newly described inflamed SCLC subtype (SCLC-I) after chemotherapy also reflects SCLC plasticity and SCLC-I predicts preferential response to immune checkpoint inhibitor combination therapy^15^. Phenotypic plasticity may explain the almost inevitable relapse and early metastatic spread as tumor cells adopt a variety of behaviors to adapt and thrive in diverse microenvironments^16^ and thus strategies to combat plasticity may be essential for effective treatment of SCLC patients. Although the importance of non-NE cells is recognized, their functions within SCLC tumors are not well understood.

VM is associated with hypoxia, cellular plasticity and metastasis in several cancer types^17-20^. We previously reported that VM occurs in SCLC, is associated with worse patient prognosis, and in a xenograft model was associated with chemoresistance and faster growth^21^. Here, in 25 CDX and 2 GEMMs^22,23^ we show that non-NE cells are pseudo-hypoxic and transcriptionally primed for VM, we demonstrate that perfusable VM vessels are formed by non-NE cells and that NE to non-NE transition is driven by NOTCH in CDX *ex vivo* cultures. In non-NE cells on Matrigel, proteins involved in cell-cell and cell-ECM adhesion enable collagen remodeling to form hollow tubular networks, a process requiring integrin β1. These data suggest that NE and non-NE cells must be targeted to combat VM-supported tumor growth and metastasis.

## Results

### CDX and GEMMs are tractable models to study VM

VM vessels were scored using periodic acid-Schiff (PAS+)/CD31-immunohistochemistry (IHC) in 25 CDX models^8^ (Figures 1A, 1D, Supplementary Table 1), including the previously unpublished CDX21 (Supplementary Figure 1) and in the RBL2 (*Trp53*^fl/fl^/*Rb1*^fl/fl^/*Rbl2*^fl/fl^) and RPM (*Trp53*^fl/fl^/*Rb1*^*fl*/fl^/*Myc*^*LSL/LSL*^) GEMMs^22,23^ (Figure 1B). VM vessels had lumens that frequently contained red blood cells (e.g., Figures 1A, 1B black arrows), suggesting perfusion and endothelial vessel connectivity (CD31+ brown stain, e.g., Figures 1A, 1B, green arrows). Perfusion through VM vessels was also inferred by intravenous (i.v.) injection of tomato lectin into mice harboring CDX09 tumors, which labels glycoproteins lining the inside of functional vessels carrying blood^24,25^. Immunofluorescence for i.v. tomato lectin (pink) and CD31 (yellow) within CDX09 tumors (Figure 1C, Supplementary Figure 2) demonstrated perfused, hollow VM vessels (tomato lectin+/CD31-, pink arrow) lined by human tumor cells and perfused, hollow endothelial vessels (tomato lectin+/CD31+, yellow arrow), providing evidence that VM vessels that support blood flow are present in CDX tumors. CDX VM vessel score (% VM vessels of total VM plus endothelial vessels) ranged from 0-87% (median 5%) (Figure 1D). Only 2/25 models (CDX08, CDX29) contained no detectable VM vessels (Figure 1D). Since VM correlated with accelerated tumor growth in a SCLC xenograft^21^ we assessed whether VM vessel prevalence increased with tumor size. Twelve CDX22P tumors (with robustly quantifiable and reproducible VM) were harvested over 8-15 weeks and VM vessels evaluated in tumors ranging from 203-1135 mm^3^ (Figure 1E). PAS+/CD31-VM vessels (pink) were present in all tumors (Figure 1F) and VM vessel score positively correlated with tumor volume *in vivo* (p=0.0083) (Figure 1G) whilst host endothelial vessel density remained stable (Figure 1H). Overall, these data indicate that CDX and GEMMs are tractable models for VM studies and that VM vessel networks are composed of perfusable, tubular structures that support tumor growth.

**Figure 1.**
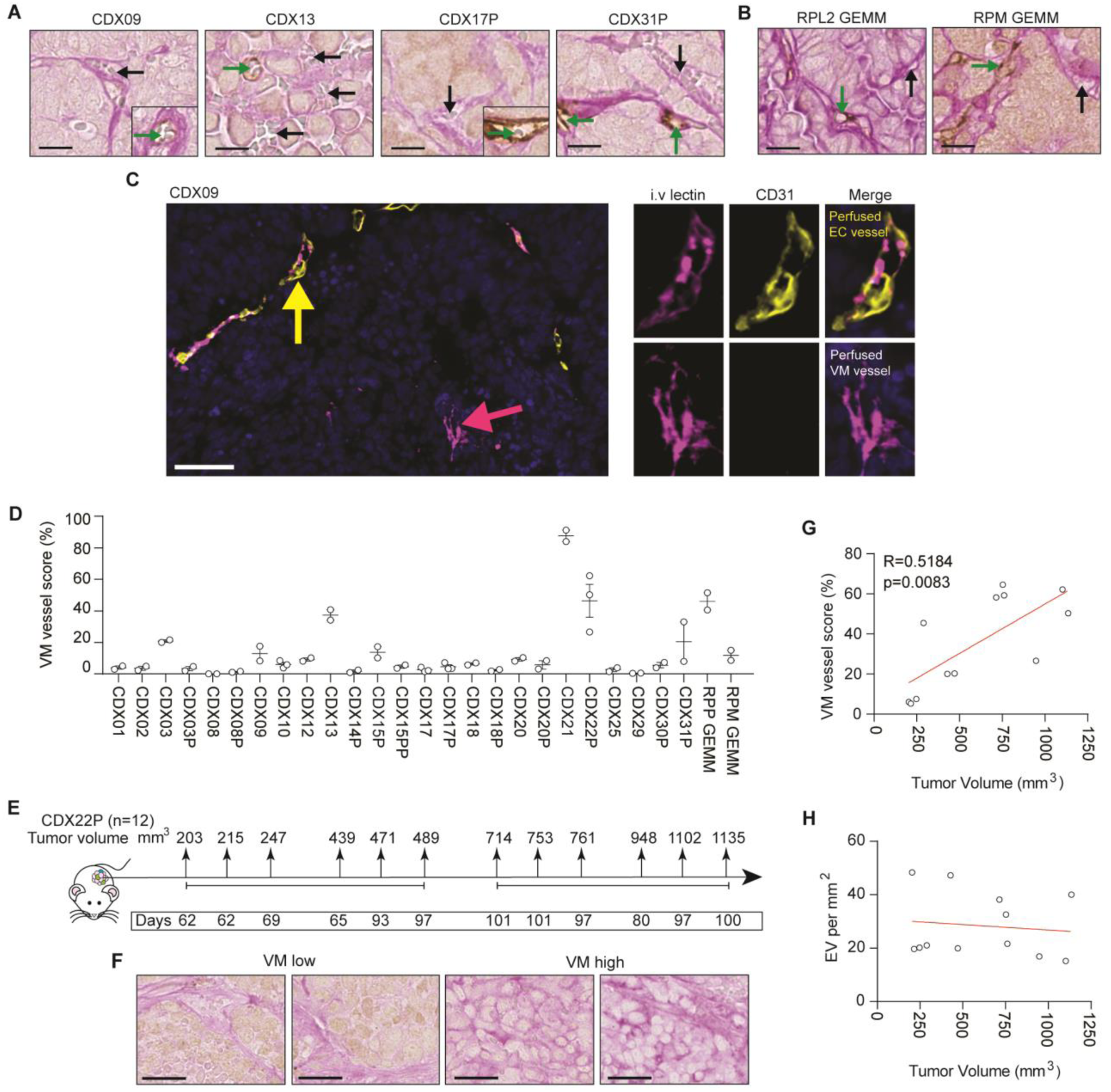
VM in SCLC CDX and GEMM. **(A)** Immunohistochemistry (IHC) of VM vessels (PAS^+^/CD31^-^, pink, black arrows) and endothelial vessels (PAS^+^/CD31^+^, brown, green arrows) in CDX. Scale bars, 200 µm. CDX models were generated from patient CTCs at pre-chemotherapy baseline and/or at post-treatment disease progression time-points (designated P). **(B)** IHC of VM vessels (PAS^+^/CD31^-^, pink, black arrows) and endothelial vessels (PAS^+^/CD31^+^, brown, green arrows) in RBL2 (*Trp53*^*fl*-/fl^/*Rb1*^fl/fl^/*Rbl2*^fl/fl^) ^22^ and RPM (*Trp53*^-fl/fl^/*Rb1fl*^/fl^/ *Myc*^*LSL*/*LSL*^) ^23^ GEMMs. Scale bars 200 µm. **(C)** Representative immunofluorescence image of a perfused endothelial (EC) vessel (CD31^+^/intravenous (i.v) tomato lectin^+^, yellow arrow) and a perfused VM vessel (CD31^-^/i.v tomato lectin^+^, pink arrow) within a CDX09 tumor that was harvested following i.v. tomato lectin injection. Single channel IF for CD31 (yellow) and i.v tomato lectin (pink) shown with merged multiplex on the right where nuclei are blue (DAPI stain). Scale bars 50 µm (left panel) and 10 µm (right panels). **(D)** VM vessel score (% VM vessels relative to total vessels (VM + endothelial) in CDX and GEMM (n = 2-3 independent tumor replicates). **(E)** Experimental design to assess VM at increasing tumor volumes. n = 12 mice were sacrificed by Schedule 1 method between 8 to 15 weeks with tumor sizes ranging 203 mm^3^ to 1135 mm^3^. **(F)** IHC of VM vessels (PAS^+^/CD31^-^, pink, black arrows) in independent CDX22P tumors ranging 203 mm^3^ to 1135 mm^3^. Scale bar 25 µm. **(G)** Pearson correlation of VM vessel score versus tumor volume. Each circle represents an independent tumor replicate. (R=0.5184, p=0.0083). **(H)** Pearson correlation of endothelial vessels (EV) per mm^2^ versus tumor volume. Each circle represents an independent tumor replicate.

### VM vessels lack NE differentiation markers and co-localize with REST^pos^ non-NE cells

Most CDX contain a majority of NE and minority of non-NE cells. We sought to determine whether one or both phenotypes were VM-competent. CDX21 contains discrete VM vessel-positive and VM vessel-negative regions (Figure 2A) for marker co-localization analysis. In serial tissue sections of CDX stained with NE markers Synaptophysin (SYP) or Neural Cell Adhesion Molecule-1 (NCAM) and the non-NE marker RE-1 Silencing Transcription Factor (REST), cells within VM-positive regions versus VM-negative regions had significantly reduced expression of SYP (15.9% versus 58.0%, p<0.0001) and NCAM (8.1% versus 39.6%, p<0.0001) and significantly higher expression of REST (40.4% versus 23.4%, p=0.002) (Figure 2B). Multiplex chromogenic IHC for VM vessels and REST (Figure 2C) or SYP (Figure 2D) in 13 CDX models, representing a range of VM vessel scores (Figure 1C) and NE to non-NE cell ratios^8^ showed that VM vessel lumens were surrounded by cells with REST-positive nuclei (Figure 2C yellow arrows) lacking SYP expression (Figure 2D). Despite the low percentage of total REST-expressing cells (<10%), the majority of VM vessels were lined with REST-positive cells (mean 74%, range 27-99%, Figure 1E). CDX13, the only non-NE POU2F3 subtype CDX^8^ was the only model with REST expressed throughout the tumor, where VM vessels were abundant (VM vessel score 38%, Figure 1C) and co-localized with REST (Figures 2C, 2E). In CDX bulk RNAseq data^8^ VM vessel score correlated positively with *REST* (Figure 2F, R=0.57, p=0.0038) and negatively with *SYP* (Figure 2G, R=-0.68, p=0.00027). In GEMM tumors, cells lining VM vessels co-localized with *REST* transcript detected by RNA *in situ* hybridization (ISH) (absolute quantification of co-localization was not feasible by ISH) (Supplementary Figure 3). Together, these co-localization data demonstrate mutual exclusivity between VM vessels and NE cells and identify the minority non-NE cell subpopulation as VM-competent in human and mouse SCLC tumors.

**Figure 2.**
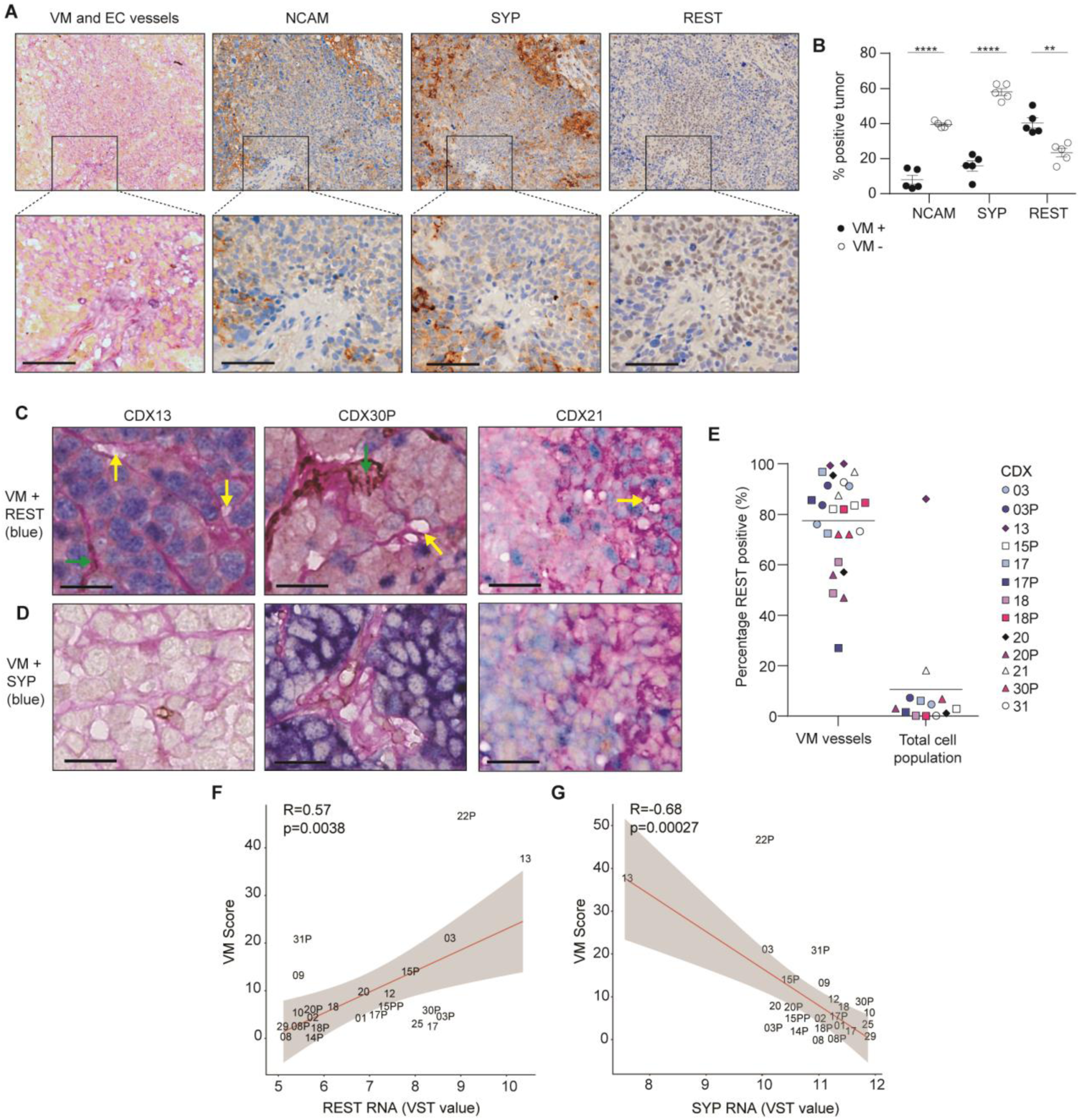
VM vessels co-localize with REST^+^ non-NE cells. **(A)** Immunohistochemistry (IHC) in serial CDX21 tissue sections for VM vessels (PAS^+^/CD31^-^, pink), endothelial vessels (PAS^+^/CD31^+^, pink+brown), REST, SYP and NCAM (brown). Scale bars, 50 µm. **(B)** Quantification of SYP, NCAM and REST expression in VM positive (black circles) and VM negative (open circles) regions of CDX identified in (**A**). Each circle represents a region that was defined and quantified, 2 independent mice were analyzed. *n* = 2 mice, *n* = 5 regions. Data are mean ± S.E.M. **p < 0.01, ****p < 0.0001 two-tailed unpaired student’s t-test. **(C, D)** Multiplex IHC showing VM vessels (PAS^+^/CD31^-^, yellow arrows), endothelial vessels (PAS^+^/CD31^+^, brown, green arrows) and REST (blue, in **C**) or SYP (blue, in **D**). Scale bars, 100 µm. **(E)** Percentage of VM vessels co-localized with REST (REST^+^ VM vessels). *n* = 2 mice per CDX. Total REST expression for each CDX was derived from^8^. Tumor sections from *n* = 3 independent mice per CDX were analyzed and the mean REST expression plotted **(F)** Pearson correlation of VM score versus *REST* transcript expression in CDX (R=0.57, p=0.0038). Average *REST* expression is shown for 3 independent tumors per CDX^8^. **(G)** Pearson correlation of VM score versus *SYP* transcript expression in CDX (R=0.68, p=0.00027). Average *SYP* expression is shown for 3 independent tumors per CDX^8^.

### NOTCH signaling drives a NE to non-NE transition that enables network formation consisting of hollow tubules in CDX *ex vivo* cultures

*Ex vivo* cultures of SCLC CDX recapitulate the phenotypic and molecular heterogeneity of CDX *in vivo*^26^, containing mixtures of suspension (NE) and adherent (non-NE) cells^27^. Separated adherent and suspension cultures were established from four CDX and their respective NE and non-NE phenotypes confirmed by marker expression (Figure 3A). CDX31P is an ASCL1 subtype whereas CDX17/17P and CDX30P are ATOH1 subtypes (no ATOH1 antibody is available) and CDX17 and CDX30P NE cells co-express NEUROD1^8^. In all models, suspension cells express SYP and adherent cells express the non-NE markers REST and YAP1^7,27^ (Figure 3A). Active NOTCH promotes NE to non-NE phenotype transition in RBL2 and RPM SCLC GEMMs^9,11^ and adherent CDX cells expressed cleaved (active) NOTCH (NOTCH1 and/or NOTCH2) (Figure 3A). Adherent cells from all four CDX also expressed MYC, which is associated with NE-low and non-NE SCLC^10,23^. Collectively, these data validate physical separation to interrogate VM in NE and non-NE subpopulations.

**Figure 3.**
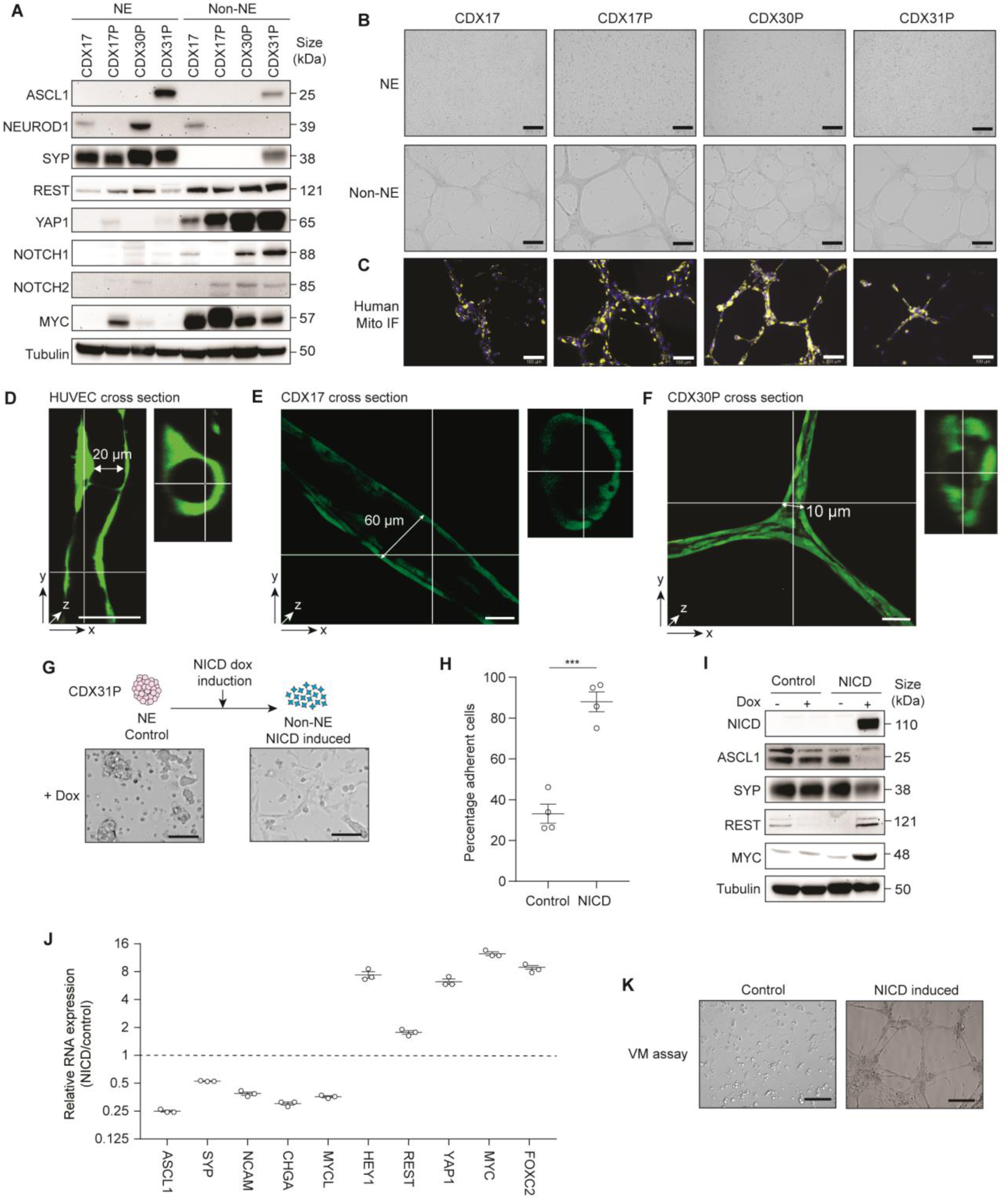
Non-NE CDX cells *ex vivo* are VM competent and require NOTCH signaling. **(A)** Representative immunoblots of CDX NE and non-NE cell lysates. *n* = 2-3 independent replicate tumors per CDX. Tubulin loading control and was run subsequently for each marker shown on the same blot **(B)** Representative brightfield images of tubule-forming assay with CDX NE and non-NE cells lysates. *n* = 2-3 independent replicate tumors per CDX. Scale bars, 500 µm. **(C)** Representative immunofluorescence of CDX non-NE cells in tubule-forming assay stained for human mitochondria (yellow) and nuclear DAPI (blue) in (B). *n* = 2-3 independent replicate tumors per CDX. Scale bars, 100 µm. **(D-F)** Representative images of Human umbilical vein endothelial cells (HUVECs) (D), CDX17 (E) and CDX30P (F) non-NE cells labelled with Cell Tracker Green forming hollow tubules when grown on Matrigel for 72 hours. Confocal microscopy images are shown after Z-stack software reconstruction (Imaris). Tubule length and diameter dimensions are shown, scale bars 50 µm. **(G)** Representative images of empty-vector control and NOTCH intracellular domain (NICD) expressing CDX31P cells three weeks post-doxycycline (dox) dox induction. Scale bars, 250 µm. **(H)** Percentage of adherent cells in control versus NICD expressing CDX31P cells three weeks post-dox induction in the suspension cells. *n* = 4 replicates from one tumor sample. Data are mean ± S.E.M. ***p < 0.001 two-tailed unpaired student’s t-test. **(I)** Representative immunoblots in control and NICD CDX31P cells ± dox. Tubulin loading control and was run subsequently for each marker shown on the same blot **(J)** Real-time quantitative PCR analysis of NE (ASCL1, SYP, NCAM, CHGA, MYCL) and non-NE (HEY1, REST, YAP1, FOXC2, MYC) markers in control versus NICD expressing CDX31P cells. Mean values are shown (black lines) where each circle represents one independent analysis, error bars are ± S.E.M **(K)** Representative images of tubule-forming assay with control and NICD CDX31P cells three weeks post-dox induction. Scale bars, 200 µm. *n* = 3 independent replicate tumors.

Formation of branching networks by cells on Matrigel is an established *in vitro* surrogate assay for VM competence^17,18,21,28^. When NE and non-NE cell subpopulations from these four CDX models were cultured on Matrigel and stained for human mitochondria, only the non-NE adherent human cells formed branching networks (Figures 3B, 3C). Similarly, only non-NE cells from both RPM and RBL2 GEMMs formed networks on Matrigel (Supplementary Figures 4A, 4B). HUVEC cells are the archetypal endothelial cell line used to study vasculogenesis and form hollow networks *in vitro* on Matrigel^29,30^ (e.g., Figure 3D). When cultured under identical conditions on Matrigel, we asked whether branching networks formed by SCLC models were comparable to the hollow tubules formed by HUVECs, thus inferring functional similarities *in vivo*. To interrogate this, fluorescently labelled CDX non-NE cells were cultured on Matrigel for three days and analyzed by fluorescence confocal microscopy. 3D reconstruction of confocal microscopy z-stacks show CDX cells form 3D tubules containing a hollow lumen (Figures 3E, 3F, Supplementary Figures 4C, 4D, where representative images are shown). Tubule length and lumen diameter varied between the three CDX models tested (CDX17, CDX17P, CDX30P) and the average tubule length was 370 μm (range 200-570 μm) with an average lumen diameter of 28 μm (range 10-60 μm) compared to 17 μm (range 14-20 μm) for HUVEC tubules. The CDX tubule diameters were greater than that of a small capillary (3-8 µm in diameter) and would be sufficient to enable flow of erythrocytes *in vivo* (∼6-8 µm in diameter)^31^.

NE to non-NE phenotype switching in SCLC GEMMs is driven by NOTCH signaling^9-11^ so we next asked whether NOTCH activation promotes NE to non-NE switching in CDX to enable VM network formation *ex vivo*. We generated CDX31P suspension NE cells with a doxycycline-inducible NOTCH intracellular domain (NICD) to drive NOTCH signaling and assessed NE to non-NE transition three weeks after NICD induction by quantification of adherent cells. After inducible expression of NICD, 88% of NOTCH-active CDX31P cells became adherent compared to 33% of empty-vector control cells (Figures 3G, 3H, P=0.0002) supporting tumor plasticity rather than solely the pre-existence of non-NE cells in CDX tumors giving rise to non-NE progeny cells. Upon NOTCH activation, NE marker expression (ASCL1, SYP) was reduced with concomitantly increased non-NE marker expression (REST, MYC) (Figure 3I). Expression (by RT-qPCR) of a larger panel of NE (*ASCL1, SYP, NCAM, CHGA, MYCL*) and non-NE (*REST, MYC, HEY1, YAP1*) markers demonstrated reciprocal expression in control versus induced NOTCH-active cells (Figure 3J). The TF *FOXC2*, recently reported as a driver of VM in multiple cancer types (Cannell *et al*., under revision) was also up-regulated 9-fold in NOTCH-active cells (Figure 3J). When cultured on Matrigel, only NICD-expressing CDX31P cells formed networks whereas control cells did not (Figure 3K). These data confirm that NOTCH signaling can drive NE to non-NE transition in a human ASCL1 subtype CDX model and that only non-NE cells are VM-competent.

### Non-NE cells express hypoxic, vascular endothelial and cell-ECM remodeling gene signatures and are transcriptionally primed for VM

To define molecular processes in SCLC VM, we profiled gene expression by RNAseq in separated NE and non-NE cells from four CDX cultured on plastic or on Matrigel (Figure 4A) and tested the hypothesis that a VM-specific signature would be observed only in non-NE cells forming networks on Matrigel. Principal component analysis (PCA) demonstrated greatest variance (29%) between NE and non-NE cell subpopulations, followed by 19% variance between CDX models (Figure 4B). As expected, suspension cell gene expression profiles aligned closely with a published SCLC NE gene signature^10^ (Supplementary Figure 5) and in non-NE cells there was increased transcription of genes associated with NOTCH pathway activation and downregulated inhibitory NOTCH pathway ligands (Supplementary Table 2). However, counter to our hypothesis, the clear phenotypic and functional differences between non-NE cells on Matrigel forming networks and those on plastic, incapable of network formation, were not reflected in differential gene expression. This implies that NE to non-NE transition transcriptionally primes cells for VM, but that additional stimuli are required to form vessels. Gene set enrichment analysis (GSEA) of NE and non-NE cells from the four CDX models revealed non-NE cell transcriptomes were enriched for blood vessel development, ECM organization and cell migration (Figure 4C) consistent with network formation on Matrigel and endothelial cell behaviors (Figures 1, 3). This is substantiated by the relative up-regulation of an endothelial-specific gene set^32^ that we have refined to remove mesenchymal genes used to implicate epithelial to mesenchymal transition^33^ in non-NE vs NE cells (Figure 4D, Supplementary Table 3), also in keeping with VM-competent cells in breast and other tumor types (Cannell *et al*., under revision), and the up-regulation of functional vascular pro-tubulogenic genes (e.g. *VEGFC, FLT1, ESM1, TIE1, TEK, CD34*) and blood coagulation cascade genes (e.g. *TFPI, TFPI2, THBD, SERPINE1/2, PLAU*) in the non-NE cells (Supplementary Table 3). Physiological hypoxia, a primary driver of angiogenesis^34^ stimulates VM in several cancer types^20,35,36^. Whilst non-NE CDX cells harbor hypoxia gene signatures (Figures 4E, 4F)^37,38^, network formation occurs in well oxygenated Matrigel. We reasoned that this paradox might be explained if non-NE cells acquire pseudohypoxic attributes^39^ and confirmed that non-NE cells (in 21% O_2_) exhibit stabilized HIF-1α and up-regulation of its downstream effectors GLUT1 and/or CA9 compared to their NE counterpart cells (Figure 4G).

**Figure 4.**
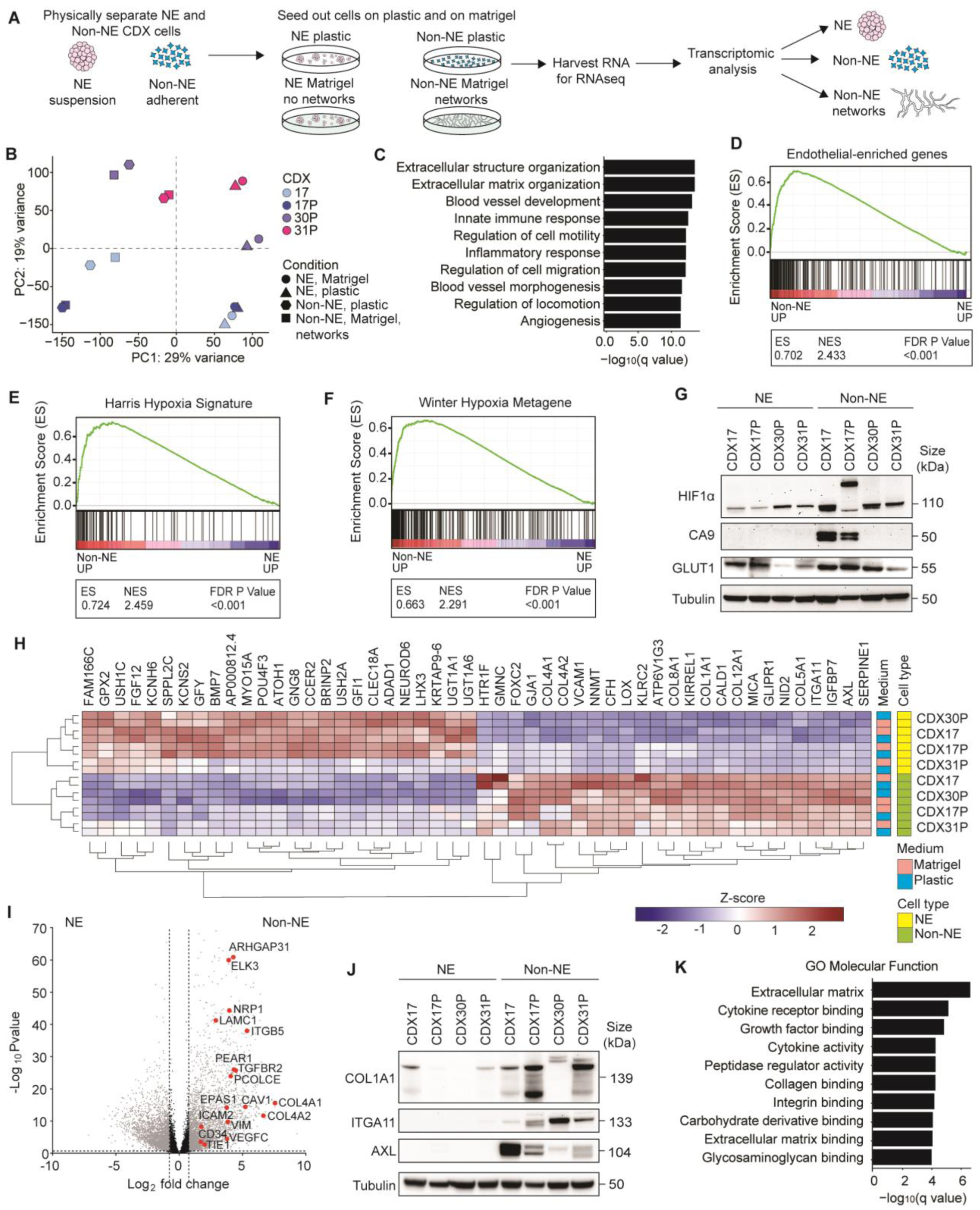
Transcriptomic analysis of network-forming CDX cells. **(A)** Workflow for generation of CDX suspension NE and adherent non-NE cells that were physically separated and cultured on plastic and Matrigel. From these samples RNA was isolated for RNA sequencing (RNAseq) followed by transcriptomic analysis. **(B)** Principal component analysis of CDX (CDX17, green, CDX17P, orange, CDX30P, purple, CDX31P, pink) NE and non-NE cell transcriptomes cultured on plastic and on Matrigel. Each symbol represents an individual replicate: circles, NE cells on Matrigel, triangles, NE cells on plastic, hexagons, non-NE cells on plastic, squares, non-NE cells on Matrigel (network forming) per CDX model. **(C)** Gene Set Enrichment Analysis (GSEA) of Biological Processes up-regulated in CDX non-NE cells compared to CDX NE cells. **(D)** GSEA of endothelial enriched genes^32^ in CDX NE and non-NE cell transcriptomes. The endothelial gene set was refined to remove any genes that are expressed in mesenchymal cells^33^. (**E)** GSEA of Harris hypoxia signature^38^ in CDX NE and non-NE cell transcriptomes. (**F)** GSEA of Winter hypoxia metagene signature^37^ in CDX NE and non-NE cell transcriptomes. **(G)** Representative immunoblots in CDX NE and non-NE cell lysates. *n* = 2-3 independent replicate tumors per CDX. Tubulin loading control and was run subsequently for each marker shown on the same blot **(H)** Heatmap of the top 25 upregulated and downregulated genes in CDX non-NE cells (green) compared to NE cells (yellow), cultured on either plastic (blue) or Matrigel (pink) **(I)** Volcano plot of differentially expressed genes in CDX non-NE cells compared to NE cells. Significant (fold change >1, -log(qvalue) >1) genes in red. **(J)** Representative immunoblots of CDX NE and non-NE cell lysates. *n* = 2-3 replicate tumors per CDX. Tubulin loading control and was run subsequently for each marker shown on the same blot **(K)** Gene Ontology (GO) Molecular Functions upregulated in CDX non-NE cells compared to CDX NE cells. For GSEA in **C, D, E, F** and **K**, NE and non-NE cells grown on plastic and Matrigel were combined and treated as technical replicates since there was no significant transcriptomic changes identified between these culture conditions.

The top 25 upregulated genes (>80 fold) in CDX non-NE cells included cell-cell adhesion receptor vascular cell adhesion molecule-1 (*VCAM1*), cell-ECM adhesion receptor *ITGA11* and multiple fibrillar and basement membrane collagen genes (*COL1A1, COL4A1, COL4A2, COL8A1, COL5A1* and *COL12A1*) (Figure 4H). The VM-associated anticoagulant^28^ and SCLC brain metastasis colonization factor^40^ *SERPINE1*, the angiogenesis associated *AXL* receptor tyrosine kinase^41^, master-regulator of VM *FOXC2* (Cannell *et al*., under revision) and endothelial-associated genes^32^ were all significantly upregulated in non-NE compared to NE cells (Figures 4H, 4I). Increased expression of COL1A1, ITGA11 and AXL proteins in non-NE CDX cells compared to their NE counterparts was confirmed in *ex vivo* cultures (Figure 4J). GSEA analysis also implicated processes associated with ECM production, collagen binding and integrin signaling in the non-NE cells (Figure 4K). Carbohydrate and glycosaminoglycan binding molecular functions were enriched in non-NE cells (Figure 4K), supporting the histopathology of VM vessels *in vivo* that are lined by PAS-positive glycoprotein basement membrane and marked by lectin when i.v injected (Figure 1). Collectively, our RNAseq data demonstrated that non-NE CDX cells are transcriptionally primed for VM given a conducive microenvironment and implicate cell-cell and cell-ECM interactions as VM-enabling processes.

### Changes in the cell adhesion proteome during non-NE cell VM

Given the functional changes that occur during network formation without obvious alteration in gene expression in non-NE CDX cells on Matrigel versus plastic, we performed a proteomic analysis of NE and non-NE cells on both substrates to identify VM-specific changes and with acellular Matrigel incorporated as an experimental control (Figure 5A, workflow schematic). We chose the RBL2 GEMM for this analysis to minimize patient-to-patient variability and exploit the ability to purify tumor cell subpopulations via FACS of NE (GFP^-^) and non-NE (GFP^+^) cells^11^. Differential protein expression analysis identified 332 significantly up-regulated proteins specific to network-forming non-NE cells on Matrigel (Figure 5B, red circles fold change>1, -log(pvalue)>1) compared to non-NE cells on plastic or NE cells on Matrigel where networks were not formed and where Matrigel components were excluded.

**Figure 5.**
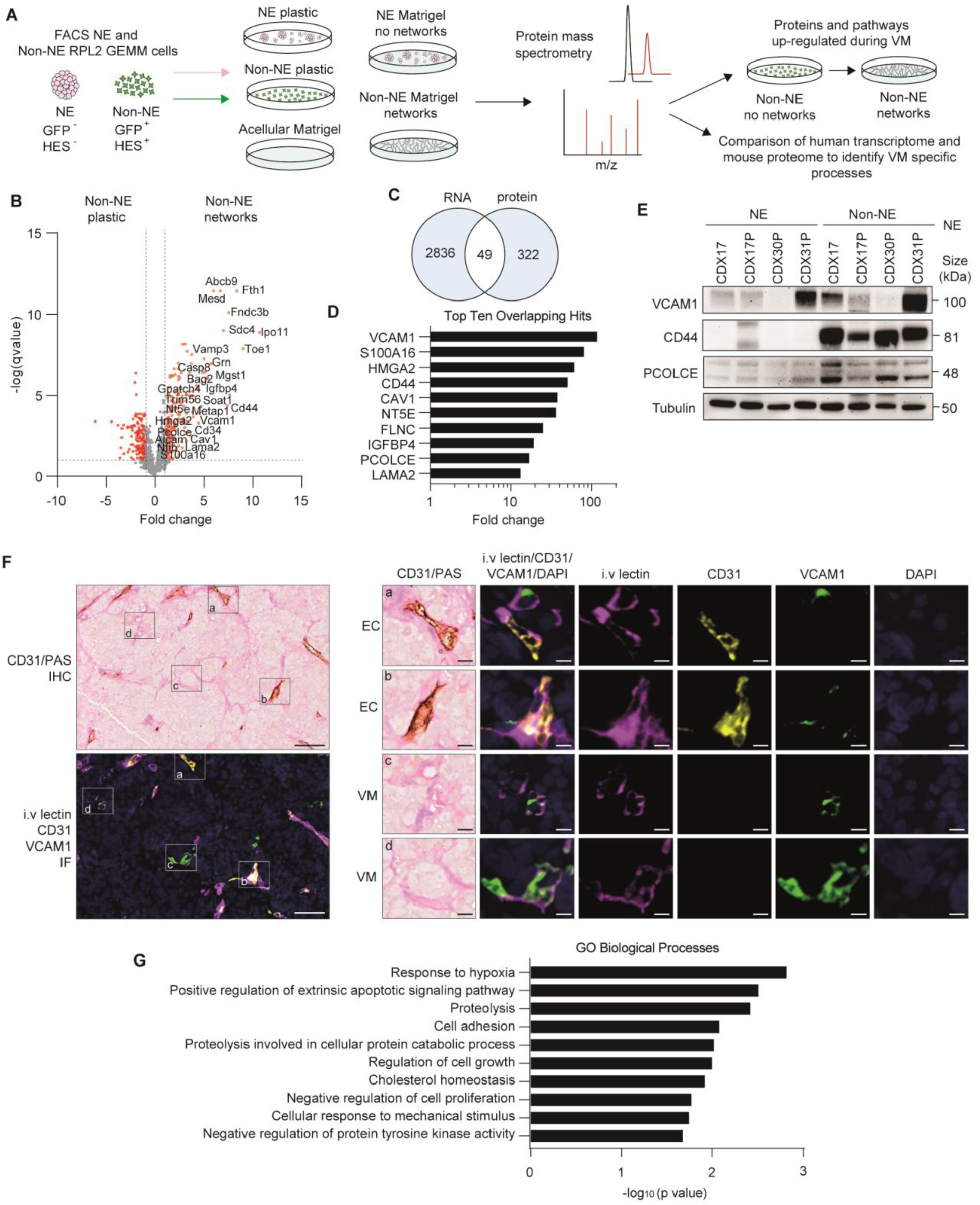
Proteomic analysis of network-forming GEMM cells. **(A)** LC-MS/MS experimental outline. NE (HES1^-^/GFP^-^) and non-NE (HES1^+^/GFP^+^) cells were generated from RBL2 GEMM tumors by flow cytometry based on *Hes1*-GFP reporter expression and seeded onto plastic and Matrigel with a blank Matrigel control. Protein lysates were harvested, processed and analyzed by LC-MS-MS in biological triplicate and technical duplicate. **(B)** Volcano plot of proteins in GEMM non-NE cells forming tubules on Matrigel versus growth on plastic. Red circles, significantly differentially expressed proteins (fold change >1, -log(pvalue) >1). *n* = 2 independent tumor replicates analyzed in triplicate per condition. **(C)** Venn diagram showing 49 VM candidates overlapping between the 322 up-regulated proteins in the network forming non-NE GEMM cells and the 2836 up-regulated genes in the CDX non-NE cells. **(D)** Transcript fold change in CDX non-NE versus NE cells of the top ten overlapping protein and RNA hits identified in (**C**). **(E)** Representative immunoblots of CDX NE and non-NE cell lysates. *n* = 2-3 replicates per CDX tumor. Tubulin loading control and was run subsequently for each marker shown on the same blot **(F)** Representative images of CD31/PAS immunohistochemistry (top left) and intravenous (i.v) tomato lectin/CD31/VCAM1/DAPI multiplex immunofluorescence (IF) (bottom left) in serial tissue sections of a CDX17P tumor harvested after mice received i.v. tomato lectin injection. Individual perfused VCAM1^+^ endothelial (EC) vessels (images a and b; i.v lectin^+^/CD31^+^/VCAM1^+^) and perfused VCAM1^+^ VM vessels (images c and d; i.v. lectin^+^/CD31^-^ /VCAM1^+^) are shown on the right with single channel IF for CD31 (yellow), i.v tomato lectin (pink), VCAM1 (green) and nuclei (DAPI, blue). Scale bars 50 µm (left panels) and 10 µm (right panels). **(G)** Gene Ontology Biological Processes representing the 49 overlapping protein and RNA hits identified in the GEMM and CDX.

Amongst the most significantly up-regulated VM-associated proteins were the cell-ECM adhesion receptor syndecan-4 (SDC4), the endothelial cell surface receptor CD34, cell-cell adhesion receptors VCAM1 and CD44, and a regulator of SCLC metastasis, Nuclear Factor I B (NFIB)^42^ (Figure 5B). We next compared these 322 up-regulated proteins with significantly up-regulated genes (p<0.05, fold change >1) in the non-NE CDX cells identified by our un-biased transcriptomics analysis on CDX models (Figure 4I) and identified 49 overlapping VM-specific candidates (Figure 5C). The top ten overlapping protein and RNA hits (Figure 5D) were up-regulated >13-fold at the transcript level in non-NE compared to NE CDX cells, again identifying VCAM1 and CD44, and procollagen processing enhancer pro-collagen C endopeptidase enhancer (PCOLCE) (Figure 5D), which were confirmed at the protein level (Figure 5E). Using multiplex IF in CDX17P tumors harvested from mice after i.v lectin injection, we showed that the endothelial marker VCAM1 was expressed within perfused endothelial vessels (Figure 5F, panels a and b: i.v lectin^+^/CD31+/VCAM1+) and perfused VM vessels (Figure 5F, panels c and d: i.v lectin^+^/CD31^-^/VCAM1+). The i.v lectin+/CD31-/VCAM1+ perfused VM vessels co-localised with PAS^+^/CD31^-^ VM vessels in an adjacent tissue section by IHC (Figure 5F) and combined, these data infer endothelial properties of VM vessels *in vivo*. Up-regulation of cell adhesion, cellular responses to mechanical stimuli and hypoxia responses was revealed by gene ontology (GO) analysis of the 49 overlapping upregulated and VM associated non-NE cell proteins (Figure 5G). Overall, the combined transcriptomic and proteomic analyses signpost the importance of cell-cell and cell-ECM adhesion and ECM remodeling processes in SCLC VM (Figures 4C, 5G).

### Integrin β1 is required for collagen remodeling *in vitro* during network formation

We hypothesized that the cell-ECM receptor integrin β1 might also be required for SCLC VM as we identified integrin signaling and ECM binding within our transcriptomics dataset (Figure 4K), and cell-ECM adhesion within our proteomics dataset (Figure 5B). Cell surface area (CSA) is a surrogate of integrin-ECM binding following short-term incubation with specific ECM substrates. CDX NE cells spread poorly both on plastic (CSA = 109 µm^2^, range 39-382 µm^2^) and on collagen (CSA = 109 µm^2^, range 45-310 µm^2^) whereas non-NE cells adhered partially on plastic (CSA = 304 µm^2^, range 87-972 µm^2^) but exhibited significantly increased spreading on collagen (CSA = 1478 µm^2^, range 135-9122 µm^2^, p<0.0001) (Figure 6A). Non-NE cell spreading on laminin was comparable to plastic (Supplementary Figure 6A), indicating a preferential collagen-mediated interaction. These data were recapitulated in three RBL2 GEMM cell lines derived from different mice (p<0.0001) (Supplementary Figure 6B). Perturbation of integrin-collagen interactions with an integrin β1 blocking antibody abrogated non-NE cell spreading on collagen (CSA = 429 µm^2^, range 137-3173 µm^2^, versus CSA = 1492 µm^2^, range 234-7119 µm2, p<0.0001) (Figure 6A) and impaired network formation *in vitro* in four CDX non-NE cell cultures (Figure 6B), where tubule branching length was significantly reduced compared to isotype control (4150 μm versus 10,461μm, p<0.0001, Figure 6C). Cytoskeletal changes transduced via FAK activation downstream of integrin β1-ECM binding^43^ were also implicated in VM network formation as integrin β1 blockade reduced p-FAK in 3 of 4 CDX non-NE cell cultures (Figure 6D).

**Figure 6.**
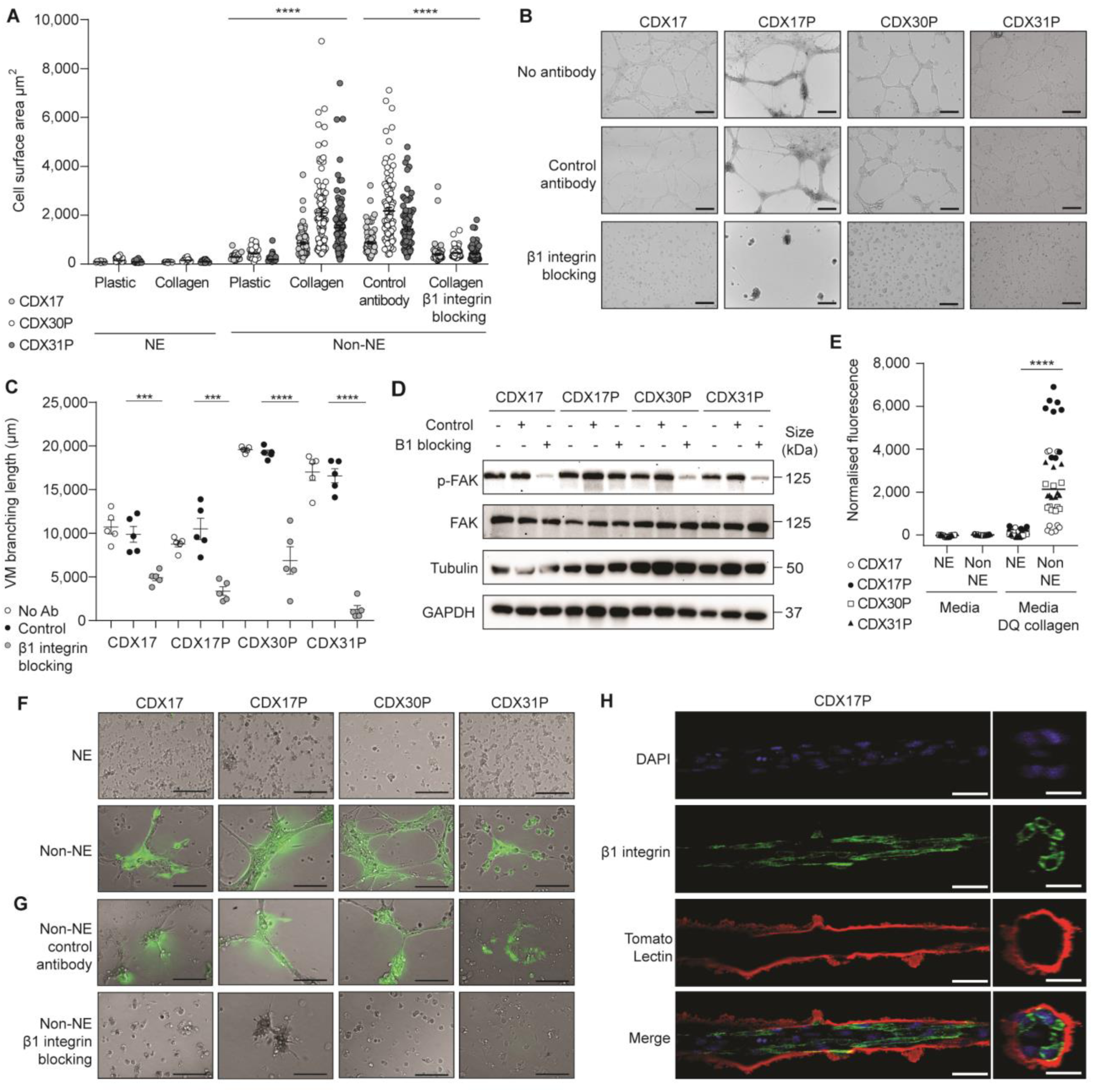
Integrin β1 is required for collagen remodeling *in vitro* during network formation. **(A)** Cell surface area (CSA) of CDX (light gray circles, CDX17, open circles, CDX30P, gray circles, CDX31P) NE and non-NE cells on plastic or collagen, and treated with an integrin β1 blocking antibody or isotype control antibody. Data are mean ± S.E.M. 200 cells analyzed from a CDX tumor. ****p < 0.0001 two-tailed unpaired student’s t-test. **(B)** Representative images of CDX non-NE cell tubule-forming assay with no antibody, isotype control antibody and integrin β1 blocking antibody. Scale bars, 500 µm. *n* = 3 replicates per CDX tumor. **(C)** VM branching length of tubule-forming assays in (**C**). Data are mean ± S.E.M, 5 images analyzed per experiment (open circles, no antibody, black circles, isotype, gray circles, integrin β1 blocking antibody), representative of *n* = 2-3 replicates per CDX tumor. ***p < 0.001, ****p < 0.0001 two-tailed unpaired student’s t-test. **(D)** Representative immunoblots of CDX non-NE cell lysates treated with no antibody, isotype control antibody or integrin β1 blocking antibody. *n* = 3 replicates per CDX tumor. Each lysate was probed separately for p-FAK with Tubulin as a loading control and total FAK with GAPDH as a loading control **(E)** Fluorescence of CDX (open circles, CDX17, black circles, CDX17P, open square, CDX30P, black triangle, CDX31P) NE and non-NE cells cultured in media ± Dye-Quenched (DQ)-collagen. Each circle represents fluorescence of a single well. *n* = 9 replicates per CDX tumor. Data are mean ± S.E.M. ****p < 0.0001 two-tailed unpaired Student’s t-test. **(F)** Representative fluorescence images of tubule-forming assay with CDX NE and non-NE cells on Matrigel containing DQ-collagen. Scale bars, 500 µm. n = 2 replicates per CDX tumor. **(G)** Representative fluorescence images of CDX non-NE cell tubule-forming assay with no antibody, isotype control antibody and integrin β1 blocking antibody on Matrigel containing DQ-collagen. Scale bars, 500 µm. *n* = 2 replicates per CDX tumor. **(H)** Representative confocal microscopy images of CDX17P non-NE cells on Matrigel with immunofluorescent staining for DAPI nuclear stain (blue), membranous integrin β1 (green) and coated by extracellular matrix containing lectin (tomato lectin, stained red). Images are shown following z-stack reconstruction using Imaris software after 72 hours on Matrigel, scale bar 50 µm.

Since non-NE cells require integrin β1 to adhere to collagen-rich ECM in Matrigel and in microvascular basement membranes^44^ we hypothesized that non-NE cells remodel collagen into VM structures both on Matrigel and in CDX tumors, identified by PAS-lined VM vessels (Figure 1). As a surrogate for collagen remodeling *in vivo*, we measured collagen cleavage by culturing CDX non-NE and NE cells in the presence of dye-quenched (DQ)-collagen (20 µg/mL) which fluoresces when cleaved. Non-NE cells cleaved significantly more (45-fold, p<0.0001) DQ-collagen than NE cells (Figure 6E) and fluorescent non-NE cell tubules reflected incorporation of DQ-collagen (Figure 6F). Inhibition of tubule formation via integrin β1 blockade diminished DQ-collagen incorporation into non-NE cells suggesting collagen remodeling is required for network formation (Figure 6G). Picrosirius Red (PSR) staining visualizes collagen fiber organization in cells and tissues and confirmed that collagen was remodeled by non-NE CDX cells forming networks on Matrigel (Figure S6C). PAS glycoprotein stain that detects basement membranes and VM vessels in tissues was also confined to regions containing non-NE cells within networks (Supplementary Figures 6D, 6E). Since we showed that hollow tubule formation by CDX non-NE cells on Matrigel (Figures 3E and 3F) occurs via integrin β1-mediated remodeling of ECM glycoproteins (Figure 6G and Supplementary Figure 6D), we visualized tubule forming CDX non-NE cells which had been fluorescently labelled with DAPI nuclear stain (blue), integrin β1 (green) and tomato lectin (red), the latter of which recognizes poly-*N*-acetyl lactosamine-type oligosaccharide moieties present in ECM^45^ (Figure 6H). The immunofluorescence showed that integrin β1-expressing CDX cells formed multicellular hollow tubules that interact with an outer glycoprotein-rich Matrigel shell labelled by lectin that is not present within the tubule lumen (Figure 6H), further supporting a functional role of integrin β1 in hollow tubule formation by SCLC non-NE cells. Overall, these findings indicate that SCLC non-NE cells require integrin β1 to interact with collagen in the ECM which is remodeled into hollow branching networks.

## Discussion

Robust, patient-representative preclinical models of SCLC support characterization of intra- and inter-tumoral heterogeneity^8,9,14,23,46^. VM is emerging as a phenomenon exploited by aggressive cancers to maintain their supply of oxygen and nutrients to support growth and metastasis^17,18^. Having previously reported that VM correlates with poor prognosis in SCLC patients^21^, we sought to identify VM cellular machinery and explore underlying molecular mechanisms using our CDX biobank which have a range of VM profiles (Figure 1). We demonstrate here that non-NE cell subpopulations are responsible for VM in human CDX and GEMM SCLC tumors (Figures 1-3), and that VM correlates with tumor growth without increased angiogenesis (Figures 1E-H).

Cancer cells undergo lineage plasticity to adapt, survive therapy, grow and metastasize^47^. NOTCH-induced transition of NE to non-NE cells occurred in *ex vivo* cultures of an ASCL1 subtype CDX, which was required for VM (Figure 3), validating the SCLC plasticity seen in GEMMs^9-11^. Non-NE CDX cells formed hollow tubules on Matrigel with a lumen diameter supportive of erythrocyte transit that closely resembled structures formed by HUVECs under identical conditions (Figure 3). Combined with *in vivo* data showing that lectin-perfused, VCAM1-positive VM vessels lacking CD31-positive murine endothelial cells are present in CDX tumors *in vivo* (Figures 1 and 5), this provides evidence that human tumor cells form VM vessels with endothelial properties enabling them to act as functional blood vessels *in vivo*.

*Ex vivo* cultures of non-NE cells from four CDX models showed enhanced attachment to plastic, down-regulated NE markers and up-regulated non-NE markers including REST and YAP1^27^, relative to their suspension NE counterparts (Figure 3). Non-NE cells expressed at least one NOTCH receptor (NOTCH1 and/or NOTCH2), and increased MYC (Figure 3). MYC and NOTCH co-operate to drive NE to non-NE transition in SCLC RPM GEMM tumors^9^ with established roles in metastasis^11,13^, and co-operation between non-NE and NE cells is essential for metastasis in the RP (*Tp53*^fl/fl^/*Rb1*^fl/fl^) SCLC GEMM^13^. VM is required for metastasis in a breast cancer model^17^ and studies to examine relationships between VM, MYC, NOTCH and metastasis in SCLC CDX are underway.

A surprising result of this study was the broadly similar transcriptomes of non-NE cells on plastic (monolayers) and Matrigel (networks), suggesting that phenotypic switching from NE cells was accompanied with acquisition of a transcriptional program, poising cells for VM in a conducive microenvironment (Figure 4). Non-NE cells upregulated the hypoxia responsive TF *FOXC2* (Figure 4H), a key driver of VM and resistance to anti-angiogenic therapy in aggressive tumor subtypes (Cannell *et al;* under revision). Hypoxia promotes VM in melanoma, colorectal cancer and Ewing’s sarcoma^20,35,36^ and whilst SCLC is fast growing and typically hypoxic^48^, network formation by non-NE cells occurred in oxygenated Matrigel. However, normoxic non-NE cells expressed hypoxia gene signatures, stabilized HIF1α and upregulated its downstream effectors (Figure 4) and the hypoxia response was also identified as the top Gene Ontology Biological Process when analyzing the 49 overlapping proteomic and transcriptomic hits identified in the GEMM and CDX (Figure 5F). Therefore, unlike other cancer types, physiological hypoxia may not be essential for SCLC VM in non-NE cells because they are pseudohypoxic, whereby the non-NE cells have retained features of previous physiological hypoxia *in vivo*. Given the roles of NOTCH and MYC in SCLC NE to non-NE plasticity, it is notable that NOTCH, MYC and hypoxia signaling are also entwined in regulating endothelial angiogenesis^49,50^. Moreover, HIF-1α can bind to the NOTCH intracellular domain to augment NOTCH signaling^51^, suggestive of a forward signaling loop once non-NE cells acquire pseudohypoxic traits. HIF1-α interacts and co-operates with oncogenic MYC to enhance expression of target genes including those involved in metabolic adaptation and the Warburg effect. Further studies are now warranted to understand the intricacies and importance of pseudohypoxia in MYC and NOTCH expressing non-NE cells undergoing VM.

We show that non-NE CDX cells remodeled ECM components in Matrigel to form hollow tubular networks lined by ECM and co-localized with integrin β1 expressing cells (Figure 6H). Integrin β1 is widely expressed^52^, present in both NE and non-NE SCLC cells. Integrin β1 expression is associated with poorer prognosis in SCLC patients^53^ and metastasis in an SCLC allograft model via FAK signaling^54^. The role(s) of integrin β1 in NE and non-NE SCLC cells are unexplored. Here we demonstrate that only non-NE cells are primed with active integrin β1 to enable VM network formation on Matrigel with downstream FAK activation (Figure 6). FAK activation is also a requirement for metastatic melanoma VM^18^. The remodeling of collagen in Matrigel by non-NE cells (Figure 6) was consistent with up-regulation of PCOLCE (Figures 4, 5), a protein that binds to and enhances the activity of collagen C-proteinase to enable fibrillar collagen formation^55^. Collagen architecture regulates cancer cell motility, invasiveness and metastasis and collagen-rich tumor regions, akin to VM vessel-dense regions identified in CDX (Figure 1), are associated with aggressive cell phenotypes^56^. Furthermore, VM-forming breast cancer cells are exquisitely sensitive to changes in collagen organization^28^, further supporting the importance of integrin-mediated ECM remodeling in VM.

In summary, we report a new aspect of SCLC intra-tumoral functional heterogeneity that arises from lineage plasticity when NE cells, the major cell type in most SCLC tumors, transition to a non-NE phenotype that can mimic endothelial cell behaviors via VM. Together with findings that non-NE cells are more chemoresistant and required for SCLC metastasis^11,13,27^, this study strongly recommends that therapeutic strategies should broaden beyond targeting NE biology and co-target VM competent non-NE cells.

## Methods

### Data code and availability

RNAseq data that support the findings of this study will be deposited in the EMBL-EBI database. R scripts used to process RNAseq data are available on GitHub (https://gitlab.com/cruk-mi/cdx-derived-cell-line-rna-seq-experiments). Source data are available from the corresponding author upon reasonable request. No algorithms or software were developed in this study. Software that was used in free and open source and details on acquiring them can be found in the associated references.

### Patient samples

The 25 patients described in this study had samples obtained between February 2012 and August 2016 following informed consent and according to ethically approved protocols: European Union CHEMORES FP6 Contract number LSHC-CT-2007-037665 (NHS Northwest 9 Research Ethical Committee) and The TARGET (tumor characterization to guide experimental targeted therapy) study, approved by the North-West (Preston) National Research Ethics Service in February 2015, reference 15/NW/0078. Patient metadata can be found in Simpson et al. 2020 and Extended data Figure. 1.

### CDX Generation

CDX models were generated as previously described^8^. In brief, 10mL of EDTA peripheral blood was collected from SCLC patients enrolled onto the CHEMORES study (07/H1014/96). CTCs were enriched via RosetteSep™ (#15167, Stem Cell Technologies) and subcutaneously implanted into the flank of 8-16-week-old non-obese diabetic (NOD) severe combined immunodeficient (SCID) interleukin-2 receptor γ– deficient (NSG) mice (Charles River). CDX models were generated from patients’ CTCs enriched from blood samples at pre-chemotherapy baseline and/or at post-treatment disease progression time-points (designated P, or PP)^8^.

### Disaggregation and culture of CDX

CDX tumors were grown to approximately 800 mm^3^ and the mice were sacrificed by Schedule 1 method. The tumors were removed and dissociated into single cells using Miltenyi Biotec’s tumor dissociation kit (#130-095-929) following the manufacturer’s instructions on a gentleMACS octo dissociator (#130-096-427), as previously described^8^. Single cells were incubated with anti-mouse anti-MHC1 antibody (eBioscience clone, 34-1-2s), anti-mouse anti-IgG2a+b microbeads and Dead cell removal microbead set (Miltenyi Biotecs #130-090-101) and applied to an LS column in a MidiMACS Separator for immunomagnetic depletion of mouse cells and dead cells. CDX *ex vivo* cultures were maintained in RPMI supplemented with HITES components (10nM Hydrocortisone, 0.005 mg/mL Insulin, 0.01 mg/mL Transferrin, 10 nM β-estradiol, 30nM Sodium selenite), Rock inhibitor added fresh (Selleckchem, Y27632), and 2.5% FBS added after one week at 37° and 5% CO_2_.

### Mouse SCLC models

RBL2 SCLC (*Trp53*^*fl*/fl^/*Rb1*^fl/fl^/*Rbl2*^fl/fl^) *Hes1*-GFP reporter mouse model was generated by J. Sage^22^ and independent cell lines were obtained from the Sage laboratory and designated the references YT326, YT330 and LJS1157. NE (HES1^-^/GFP^-^) and non-NE (HES1^+^/GFP^+^) cells were separated by flow cytometry based on *Hes1*-GFP reporter expression. RPM SCLC (*Trp53*^-fl/fl^/*Rb1*^*fl/fl*^/*Myc*^*LSL*/*LSL*^) mice were generated by T. G. Oliver^23^, FFPE tissue was obtained from the Oliver laboratory and the mouse model was obtained from The Jackson Laboratory, stock no. 029971.

### Mouse Lung Tumor Initiation

For *in vivo* studies with the RBL2 model, tumors were induced in 8 to 12 weeks old mice by intratracheal instillation with 4×10^7^ plaque forming units (PFU) of Adeno-CMV-Cre (Baylor College of Medicine, Houston, TX). For *in vivo* studies with the RPM model, tumors were induced in 6 to 8 weeks old mice by nasal inhalation with 10^6^-10^8^ PFU of Adeno-CGRP-Cre (Viral vector Core, University of Iowa). Viruses were administered in a Biosafety Level 2 room according to Institutional Biosafety Committee guidelines. Both male and female mice were equally divided between treatment groups for all experiments. To generate RPM cell lines, 7-week-old RPM mice were sacrificed by Schedule 1 method and the lungs disaggregated with Liberase at 37°C (Millipore Sigma, #5401127001), according to the manufacturer’s instructions. Cell lines were maintained in RPMI 10% FBS.

### Ethics statement

For *in vivo* studies with the CDX models, all procedures were carried out in accordance with Home Office Regulations (UK), the UK Coordinating Committee on Cancer Research guidelines and by approved protocols (Home Office Project license 40-3306/70-8252/P3ED48266 and Cancer Research UK Manchester Institute Animal Welfare and Ethical Review Advisory Body). For *in vivo* studies with the RBL2 model, mice were maintained according to practices prescribed by the NIH at Stanford’s Research Animal Facility (protocol #13565). Additional accreditation of Stanford animal research facilities was provided by the Association for Assessment and Accreditation of Laboratory Animal Care (AAALAC). For *in vivo* studies with the RPM model, mice were maintained according to practices prescribed by the University of Utah’s Institutional Animal Care and Use Committee.

### Plasmids and Lentiviral Production

The human NOTCH 1 intracellular domain (h1NICD) doxycycline-inducible expression plasmid (pLIX-h1NICD) was a gift from Julien Sage (Addgene #91897)^11^. The pLIX_403 vector (a gift from David Root, Addgene #41395) was used as an empty vector control. Lentiviral vectors were packaged into lentivirus particles by co-transfecting Lenti-X 293T cells (Clontech) with pMDLg/pRRE (a gift from Didier Trono, Addgene #12251), pCMV-VSV-G (a gift from Bob Weinberg, Addgene #8454) and pRSV-Rev (a gift from Didier Trono, Addgene #12253). Lentiviral particles were harvested, filtered (0.45 µm) and CDX cells were infected with 1 mL virus containing 12 µg/mL polybrene (Sigma), followed by selection with Puromycin (Merck, P8833).

### CDX Longitudinal Growth Study

A total of 100,000 viable CDX22P cells in 100 μL 1:1 RPMI: Matrigel were injected s.c into the right flank of *n* = 12 8-12-week-old female NSG mice. Mice were randomized deterministically into four groups when tumors reached 150-200 mm^3^, to be removed at 250 mm^3^, 500 mm^3^, 750 mm^3^ or 1000 mm^3^. This avoided bias of the fastest growing tumors into one group and meant that different tumor growth rates were represented in each group. *n* = 3 tumors were allocated to each of the four size groups. The study was designed to provide *n* = 12 tumors at varying sizes to calculate correlations between tumor size and the vasculature. No animals, experimental groups or data points were excluded from the study. Blinding was not performed during this experiment as it was an exploratory study and not hypothesis testing. Tumors were harvested in ice cold formalin for formalin fixed, paraffin embedded (FFPE) tissue and each tumor was analyzed as an individual biological replicate. FFPE tissue was analyzed by immunohistochemistry (IHC) for VM vessels and endothelial vessels (see below). Linear regression analysis (*n* = 12) of VM vessel score or endothelial vessel score versus tumor volume was performed.

### CDX tumor perfusion study

A total of 100,000 viable CDX cells in 100 μL 1:1 RPMI: Matrigel were injected s.c into the right flank of 8-12-week-old female NSG mice. CDX tumors were grown to approximately 750 mm^3^ and mice received an i.v injection of Biotinylated Tomato Lectin (4mg/kg, Vector Laboratories, B-1175-1) one hour before they were sacrificed by Schedule 1 method. Tumors were excised and processed to FFPE tissue or snap frozen in liquid nitrogen, followed by immunofluorescence analysis for i.v tomato lectin and endothelial vessels (see below).

### Immunohistochemistry and *in situ* hybridization

FFPE CDX tumors and GEMM tumors were cut as 4 µm sections and stained by immunohistochemistry (IHC) for markers detailed in Supplementary Table 4. All IHC was standardized on a Leica Bond Max or Rx Platform using standard protocol F with Bond Polymer Refine Detection kit (DS9800) or on a Roche Ventana Ultra with UltraMap DAB IHC Detection kit (760-151), unless otherwise stated. For VM vessel staining CD31 was automated on the Leica Bond Max using standard protocol F minus hematoxylin. Periodic Acid Schiff (PAS) was performed manually by incubation in 4 mg/mL periodic acid (Sigma Aldrich, 375810) for 5 minutes followed by incubation in Schiff’s fushin-sulfite reagent (Sigma Aldrich, S5133) for 30 minutes in the dark, before incubation in warm water for 4 minutes and rinsing in water until clear. Chromogenic detection of biotinylated i.v lectin in FPPE tissues was automated on the Leica bond Rx using standard protocol F with the post primary mouse link and secondary detection steps substituted for a streptavidin-biotin-HRP step (Vectastain Elite ABC-HRP Peroxidase kit, Vector labs, #PK-6100).

Multiplex chromogenic IHC staining was performed on the Leica Bond Max using protocol F minus hematoxylin for CD31, followed by REST or SYP using protocol J with Bond Polymer Refine Red Detection Kit (DS9390) minus hematoxylin and red parts A, B, C and D. By manual IHC, slides were then incubated with Vector Blue reagent (Vector labs, SK-5300) for 30 minutes. PAS staining was performed thereafter. ISH for mouse REST (RNAscope® LS 2.5 probe – Mm-Rest, ACDBio, #316258) was performed on the Leica Bond RX and developed with Vector Blue chromagen. VM vessel staining on the Bond Max and PAS staining was performed thereafter.

Whole sections were scanned using a Leica SCN400 or Olympus VS120 and single plex chromogenic staining was quantified using HALO. For VM vessel scoring, VM vessels and endothelial vessels were counted manually where endothelial vessels are PAS^+^/CD31^+^ and VM vessels are PAS^+^/CD31^-^ structures with a defined lumen, sometimes containing RBCs. PAS+ mouse stromal cells (defined morphologically by distinct PAS staining patterns through the tumor within regions that do not contain SCLC tumor cells) are excluded from scoring and PAS^+^ VM vessels are scored only within regions containing tumor cells. A VM vessel score was determined as the ratio of VM vessels to total vessels and expressed as a percentage. For quantification of REST positive VM vessels in multiplex chromogenic IHC, VM vessels were identified and scored as positive if the vessel lumen was surrounded by two or more REST positive cells.

### Immunoblotting

SCLC cells were lysed on ice with lysis buffer (Cell Signaling Technologies, #9803S) containing a cocktail of protease inhibitors (Sigma, #P8340) and phosphatase inhibitors (Sigma, #P0044 and #P5726). Protein concentrations were measured using the BCA Protein Assay reagent kit (ThermoFisher Scientific, #23225). 20 μg of each protein lysates were separated by SDS-PAGE on 4-12% gradient gels (NuPAGE™, ThermoFisher Scientific, #NP0322) and transferred onto PVDF membrane (ThermoFisher Scientific, #10617354). Membranes were blocked with 5% milk diluted in Tris-Buffered Saline (TBS) 0.1% Tween for 1 hour at room temperature and incubated with primary antibodies (Supplementary Table S5) overnight at 4°C, followed by incubation with goat anti-rabbit IgG HRP (Agilent Technologies/P0440801-2), rabbit anti-mouse IgG HRP (Agilent technologies/P044701-2), goat anti-rat IgG HRP (Abcam/ab57057) or rabbit anti-goat IgG HRP (Agilent technologies/P044901-2) secondary antibodies (1:5000). Western blots were developed with Western Lightning™ chemiluminescence reagent plus (Perkin Elmer, #NEL104001EA) and imaged on a ChemiDoc™ (Bio-Rad). All blots were subsequently re-probed for a sample loading control (Tubulin or GAPDH) on the same blot. *n* = 2-3 lysates from independent tumor replicates were run independently on different blots (one representative blot shown per experiment).

### *In vitro* tubule formation assay

Culture dishes were coated with Growth Factor Reduced Matrigel (Corning #354230) and incubated at 37°C for 60 minutes to set. SCLC cells were seeded onto 6 well plates at a density of 1.5 × 10^6^ cells and imaged by phase contrast microscopy after 24 hours. For integrin β1 blocking antibody experiments, cells were incubated in media containing 10 µg/mL blocking antibody (Purified Rat Anti-Human CD29 Clone Mab 13, BD Biosciences, #552828) or equivalent concentration non-targeting isotype control (Purified Rat IgG2a κ Isotype Control Clone R35-95, BD Biosciences, #553927). To quantify tubule branching length, 4-5 random field of view brightfield images were taken per well and images were analyzed in ImageJ using the angiogenesis analyzer algorithm. For confocal imaging, cells were labelled with Molecular Probes CellTracker Green CMFSA Dye (ThermoFisher Scientific, C2925, 1 µg/mL) or CellTracker Deep Red Dye (ThermoFisher Scientific, C34565, 1 µg/mL) for 30 minutes according to the manufacturer’s instructions, and 1.5 × 10^6^ labelled cells were seeded out onto a 35mm dish coated with Growth Factor Reduced Matrigel (Corning #354230) and allowed to form tubules. Tubules were imaged on a Leica TCS SP8 confocal microscope (Leica Microsystems) using a 25X water lens and Z-stacks (40-150 µm) were reconstructed using Imaris Imaging software (Oxford Instruments).

### Immunofluorescence

Cryosectioned CDX tumors with i.v tomato lectin were fixed with 4% PFA for 10 minutes, blocked with 1% BSA, 0.3 M glycine, 0.1% Triton X-100 in PBS for 30 minutes and incubated with anti-murine CD31 (1:1000, Abcam, ab124432) and anti-human mitochondria (1:250, Abcam, ab92824) for 1 hour at room temperature, followed by Goat anti-rabbit AlexaFluor-488 (1:1000, ThermoFisher Scientific, #A-11034), Goat anti-mouse AlexaFluor-647 (1:1000, ThermoFisher Scientific, #A-21235) and streptavidin-PE (1:100, Biolegend, 405203) secondary antibodies overnight at 4°C. Tissues were mounted and whole sections were scanned on an Olympus VS120 at 20X.

For multiplex IF in CDX FFPE tumors, tissues were cut as 4 μm sections and automated IF was performed on a Leica Bond Rx Platform at room temperature using the PerkinElmer Opal 4-Colour Automation IHC Kit (PerkinElmer, #NEL800001KT). Tissue sections were blocked with 3% Hydrogen peroxide (Sigma-Aldrich, H1009) for 10 minutes to block endogenous peroxidase activity, followed by 10% Casein solution (Vector Laboratories, #SP-5020) for 10 minutes to block non-specific antibody binding. Slides were stained with primary antibody (CD31 or VCAM1) followed by DAKO envision+ system horseradish peroxidase (HRP) conjugated secondary antibody (DAKO, #K4003) for 30 minutes, followed by incubation with OPAL Tyramide-fluorophore (PerkinElmer, OPAL650, OPAL570 or OPAL520 1:200) for 10 minutes. For detection of more than one epitope, tissues were heat inactivated following the Tyramide-fluorophore incubation step, then blocked and probed with another primary antibody as above and incubated with a different Tyramide-fluorophore. Biotinylated i.v lectin was detected with the Vectastain Elite ABC-HRP Peroxidase kit (Vector labs, #PK-6100) followed by an OPAL Tyramide-fluorophore. Tissues were counterstained with nuclear DAPI (0.1 μg/mL, Fisher Scientific, #10184322) for 10 minutes, slides mounted in Molecular Probes ProLong Gold Antifade and whole sections were scanned on an Olympus VS120 at 20X.

For human mitochondria immunofluorescence 8-well Millicell slides (Millipore #PEZGS0816) were coated with Growth Factor Reduced Matrigel (Corning #354230) and seeded with cells at a density of 1.5 × 10^6^ mL^-1^. After 24 hours cells were fixed with 4% PFA for 30 minutes and permeabilized in 1% BSA, 0.3 M glycine, 0.1% Triton X-100 in PBS for 1 hour. Fixed cells were incubated with anti-human mitochondria antibody (Abcam, ab92824) at 1:250 dilution overnight at 4°C and anti-mouse AlexaFluor-555 secondary antibody (1:1000, ThermoFisher Scientific, #A-21424) for 1 hour at room temperature, followed by nuclear DAPI (1:10,000). Cells were mounted and imaged by fluorescence microscopy.

For tomato lectin and integrin β1 immunofluorescence, 35 mm petri dishes coated with Growth Factor Reduced Matrigel (Corning #354230) and seeded with cells at a density of 1.5 × 10^6^ mL^-1^. After 3 days cells were fixed with 4% PFA for 30 minutes and permeabilized in 1% BSA, 0.3 M glycine, 0.1% Triton X-100 in PBS for 1 hour. Fixed cells were incubated with integrin β1-FITC (CD29-FITC, Beckman Coulter, IM0791U, 1:25) and Tomato Lectin Biotinylated (1:250, Vector Laboratories, B-1175-1) for one hour at room temperature followed by APC-Streptavidin secondary (1:500, Biolegend, 405207) and nuclear DAPI (0.1 μg/mL, Fisher Scientific, #10184322). Cells were imaged on a Leica TCS SP8 confocal microscope (Leica Microsystems) using a 25X water lens and Z-stacks (40-150 µm) were reconstructed using Imaris Imaging software (Oxford Instruments).

### Reverse transcriptase–qPCR

RNA was isolated from CDX cells using the RNeasy mini kit according to Qiagen recommendations. cDNA synthesis was performed with the High-Capacity cDNA Reverse Transcription Kit (ThermoFisher Scientific). RT–qPCR was performed using Taqman gene expression master mix and gene expression assays for *ASCL1* (Hs00269932_m1), *SYP* (Hs00300531_m1), *NCAM* (Hs00941830_m1), *CHGA* (Hs00900370_m1), *MYCL* (Hs00420495_m1), *HEY1* (Hs05047713_s1), *REST* (Hs05028212_s1), *YAP1* (Hs00902712_g1), *FOXC2* (Hs00270951_s1), *MYC* (Hs00153408_m1), *ATOH1* (Hs00944192_s1), *NEUROD1* (Hs01922995_s1) and *ACTB* (Hs01060665_g1) according to the manufacturer’s recommendations. Data were analyzed with the dCt method by normalizing to *ACTB* housekeeping gene.

### RNAseq and transcriptomic analysis

RNA was extracted from 3–6 independent replicate tumors per CDX and RNAseq was performed as previously described^8^. Transcriptomic analysis was performed with amendments to the previously described alignment (NF-core RNAseq pipeline with STAR aligner) and annotation (mapped to Ensembl version 99)^8^. CDX NE and non-NE cells were cultured on plastic and Matrigel for 24 hours, followed by RNA extraction (RNeasy Mini Kit, Qiagen, #74104) and sequencing. Data was aligned using STAR ^57^ to GRCH38 Ensembl version 99 ^58^ as part of the RNA-Seq pipeline from nf-core^59^. Aligned reads were filtered to remove mouse contamination reads using the bamcmp algorithm ^60^ before being mapped to genomic annotation^61^. Downstream analysis was performed in R^62^. Differentially expressed genes were called using DESeq2, ^63^ log2 fold change were shrunk using the ‘ashr’ transform^64^ and visualized using the ‘EnhancedVolcano’ package. Gene set enrichment analysis was performed using GAGE^65^. For visualizations, the raw counts were transformed via the variance stabilizing transform in DESeq2.

### LC-MS/MS protein preparation

Two independent GEMM RBL2 *Hes1* reporter lines were separated into NE (GFP^+^) and non-NE (GFP^-^) subpopulations and cells were cultured on plastic and Matrigel for 24 hours, followed by protein lysate preparation and protein concentration quantification (BCA assay); performed in technical triplicate. Sample volume was adjusted to 50 µL by adding more buffer or concentrating using speed vacuum. 50 µL of 8 M urea (Sigma-Aldrich) was added to each sample and protein disulphide bonds were reduced with 5 µL of 200 mM Tris(2carboxyethyl) phosphine (TCEP, Sigma-Aldrich) solution and incubated at room temperature for 1.5 hours. Reduced disulphide bonds were capped by adding 7.5 µL of 200 mM iodoacetamide (Acros Organics) solution and incubated for 45 minutes at room temperature in the dark. Following incubation, samples were de-salted and trypsin digested following a mini S-trap column protocol provided by the manufacturer (Protifi). Briefly, 100 µL of 10% SDS solution, 10 µL of 12% aqueous phosphoric acid, and 1.4 mL of binding buffer were added to the samples, in the order described in the protocol, and vortexed. Acidified lysates were loaded onto S-trap mini spin columns in three aliquots of 500 µL and centrifuged at 4,000 x g for 60 seconds, collecting the flow-through, until all the lysate had passed though. The flow-through was reloaded again as described above. S-Trap columns were washed with 400 µL binding buffer three times, transferred to a new 2.0 mL Eppendorf tube and S-Trap columns incubated with 150 µL trypsin enzyme digestion buffer (1:30 trypsin enzyme (Thermo Fisher Scientific): protein by weight) overnight at 37 °C. Tryptic peptides were eluted from the S-Trap column via centrifugation at 1,000 x g for 60 seconds and for shotgun proteomics analysis 5 µL of the tryptic digest solution from each sample was dried down using a speed vacuum and reconstituted back into solution by adding 12 µL of 0.1% formic acid in water. 3 µL injections of each sample were analyzed by LC-MS/MS in triplicate.

### LC-MS/MS Analysis

Shotgun proteomics was performed on a LTQ-Orbitrap Elite mass spectrometer (Thermo Fisher Scientific) connected to a Dionex UltiMate RS 3000 nano-LC (Thermo Fisher Scientific). Peptides were loaded onto a C18 trap column (Acclaim PepMap, 100 A 5 µm particle size, Thermo Fisher Scientific) at a flow rate of 5 µL/min in Solvent A (0.1% formic acid in water) and desalted for 10 min. Tryptic peptides were then separated by a reversed-phase C18 analytical column (25 cm long, packed with Magic AQ C18 resin (Michrom Bioresources)). Peptides were eluted by changing the concentration of Solvent B (0.1% formic acid in acetonitrile) from 2% (first 10 min), to 35% (over 100 min), and to 85% (next 2 min followed by 5 min hold at 85%). Eluted peptides were subjected to MS1 and MS/MS on the mass spectrometer. The MS1 mass resolution was set to 60,000 with a scan range of 400-1800 m/z. The top 10 most abundant ions in each MS1 scan were selected for collision-induced dissociation (CID). Dynamic exclusion was set for 30 seconds.

### Total protein raw data analysis

MS raw files from the shotgun proteomics analysis were searched against the Swiss-Prot mouse database using Byonic (Protein Metrics) software. Quantitative information was extracted from MS1 spectra of all identified peptides using an in-house R script based on MSnbase package (10.1093/bioinformatics/btr645) and integrated from the spectrum to protein level using the WSPP model (10.1021/pr4006958) with the SanXoT software package (10.1093/bioinformatics/bty815). In summary, every scan *x*_*qps*_ = log_2_ *A*/*B* was calculated using the area under the curve of the extracted ion current (XIC) coming from group 1 and group 2. The statistical weight *w*_*qps*_ of each scan was calculated as the maximum AUC of the pair of samples to compare. The log2-ratio of every peptide (*x*_*qp*_) was calculated as the weighted average of its scans, whereas the quantification of each protein (*x*_*p*_) was the weighted average of its peptides, and the grand mean 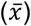 as the weighted average of all the protein measures. The variances at the scan, peptide, and protein levels, as well as protein abundant changes were determined only with non-modified peptides.

### Proteomics downstream data analysis

Global data analysis, all heatmaps, and hierarchical clustering was performed using Morpheus software (https://software.broadinstitute.org/morpheus/) from the Broad Institute. For analysis of shotgun proteomics, Matrigel contaminants were excluded and t-tests performed between the non-NE Matrigel and non-NE plastic samples. Significant differentially expression proteins were then compared to the NE Matrigel samples to generate non-NE Matrigel specific (VM, network forming) up-regulated and down-regulated proteins. Fold-change was plotted against -log(q-values) in GraphPad prism to generate volcano plots. Gene ontology analysis was performed using DAVID functional annotation tool online (https://david.ncifcrf.gov/).

### ECM adherence assays

ECM adherence assays were performed as previously described^66^. Dishes were incubated with 1 μg/ml collagen 1, 1 μg/ml laminin or PBS overnight at 4°C, followed by incubation with heat denatured BSA solution (10 mg/ml, Sigma-Aldrich, A3608) for 30 minutes, prior to rinsing in PBS and plating cells. Plates were fixed with 5% glutaraldehyde for 30 minutes and cells counterstained with 1% Crystal Violet solution in dH_2_O and imaged. Surface area of cells was determined in ImageJ by automatic cell detection following thresholding. For integrin β1 blocking antibody experiments, cells were incubated in media containing 10 µg/mL blocking antibody or equivalent concentration non-targeting isotype control, as above.

### ECM remodeling assays and *in vitro* staining

Cells on Matrigel were fixed with 4% PFA for 10 minutes and washed once in PBS. For Collagen assessment by PSR staining, cells were incubated for 1 hour at room temperature with PSR staining solution before rinsing twice in Acetic Acid solution (Picrosirius Red Stain Kit, Abcam, ab150681) and imaging. For glycoprotein assessment, cells were incubated for 5 minutes in 0.5% Periodic Acid (Sigma Aldrich, 375810), rinsing wells twice in PBS, staining with Schiff’s fushin-sulfite reagent (Sigma Aldrich, S5133) for 15 minutes, rinsing extensively in PBS, and imaging by light microscopy.

### Dye-Quenched collagen assays

To assess collagen cleavage *in vitro*, cells were incubated for 72 hours in media or media containing 20 µg/mL Dye-Quenched (DQ) collagen (Invitrogen™, DQ™ Collagen, type I from Bovine Skin, Fluorescein Conjugate, D12060) and fluorescence was measured on a plate reader and normalized to media-only containing wells with and without DQ-collagen. To assess DQ-collagen remodeling during VM formation *in vitro*, DQ-collagen was added to Matrigel at a concentration of 100 µg/mL and a VM assay performed as described above. Brightfield and fluorescence images were acquired, processed (background removal standardized to all images within the same experiment and green false colour on fluorescence images) and overlayed in ImageJ.

### Quantification and statistical analysis

Statistical tests were performed using GraphPad Prism or Excel. Error bars show mean +/- SEM unless otherwise specified. Significance was determined by the Student’s two-tailed unpaired t tests with 95% confidence intervals and p values, 0.05 considered statistically significant, unless otherwise indicated. All statistical details are further described in respective figure legends.

## Author contributions

K.L.S. and C.D. supervised and devised the study. S.M.P., K.L.S. and C.D. co-wrote the manuscript. S.M.P. carried out immunohistochemistry, CDX *ex vivo* work, immunoblotting, DQ-collagen assays and coordinated the *in vivo* study described, including data analysis and interpretation. With the exception of mass spectrometry S.C.W carried out all RBL2 GEMM work described, adhesion assays and ECM remodeling assays, including data analysis and interpretation, with help from Y.T.S. S.H carried out bioinformatics analyzes and interpretation. F.G.M and A.B carried out the mass spectrometry data acquisition that was overseen by S.J.P which was analyzed by E.H, S.M.P, F.G.M and A.B. J.H and M.H helped advise on adhesion assays and integrin blocking experiments and reviewed the manuscript. K.F. has oversight of all CDX model generation and reviewed the manuscript. M.G. is responsible for all *in vivo* work described. A.K. has oversight of all bioinformatics analysis in the Cancer Biomarker Centre. M.C, L.P and F.B. oversaw the acquisition of ethical permission and patient consent and the collection of blood samples for patients on the CHEMORES study. F.B. assisted with manuscript revision and is the chief investigator of the CHEMORES study. T.G.O supplied RPM GEMM tissue for VM analysis and contributed to data interpretation. J.S helped edit the manuscript and is responsible for all RBL2 GEMM work described. I.C carried out bioinformatic analyzes and interpretation and helped edit the manuscript with G.H.

## Disclosure of funding

This work was supported via Core Funding to CRUK Manchester Institute (grant no. C5759/A27412), Cancer Research UK Manchester Centre award [CTRQQR-2021\100010], the CRUK Lung Cancer Centre of Excellence (grant no. A20465), CRUK program grant (to M.H, grant no. C13329/A21671), NIH grant (to J.S, grant R35 CA231997; to S.P, grants U01 CA207702 and CA226051), to T.O, grants U24 CA213274 and U01 Ca231844; The Christie Charitable Fund and a CRUK-Fulbright Scholarship (to S.C.W). Support was received by the NIHR Manchester Biomedical Research Centre. Patient recruitment was supported by the NIHR Manchester Biomedical Research Centre and NIHR Manchester Clinical research Facility at The Christie Hospital. Sample collection was undertaken via the CHEMORES Trial (molecular mechanisms underlying chemotherapy resistance, therapeutic escape, efficacy and toxicity—improving knowledge of treatment resistance in patients with lung cancer).

## Declaration of interests

C.D has received research funding from AstraZeneca, Astex Pharaceuticals, Bioven, Amgen, Carrick Therapeutics, Merck AG, Taiho Oncology, Clearbridge Biomedics, Angle PLC, Menarini Diagnostics, GSK, Bayer, Boehringer Ingelheim, Roche, BMS, Novartis, Celgene, Epigene Therapeutics Inc and Thermo Fisher Scientific. C.D has received consultancy fees/honoraria and advisory board from AstraZeneca, Biocartis, Merck AG. J.S. has received research funding from Stemcentrx/Abbvie and Pfizer and licensed a patent to Forty Seven Inc/Gilead on the use of CD47 blocking strategies in SCLC. G.H and I.C have filed a patent covering use of FOXC2 and FOXC2-regulated gene sets as diagnostics and as a route toward development of VM inhibitors.

