## Supplementary Figures and Tables for "Lineage plasticity in SCLC generates non-neuroendocrine cells primed for vasculogenic mimicry"

**Supplementary Figure 1, Related to Figure 1. Clinical Features of CDX21 donor patient.**

A

| Gender | Age at Diagnosis (years) | SCLC Stage | Cell Search ® CTCs | Site(s) of Metastasis | 1st Line Treatment | No. of Cycles | Treatment Response | Survival (months) |
|---|---|---|---|---|---|---|---|---|
| Male | 80 | Extensive | 27 | Bone, Pleural Effusion, Adrenals | Carboplatin | 1 | Stable response | 2.1 |

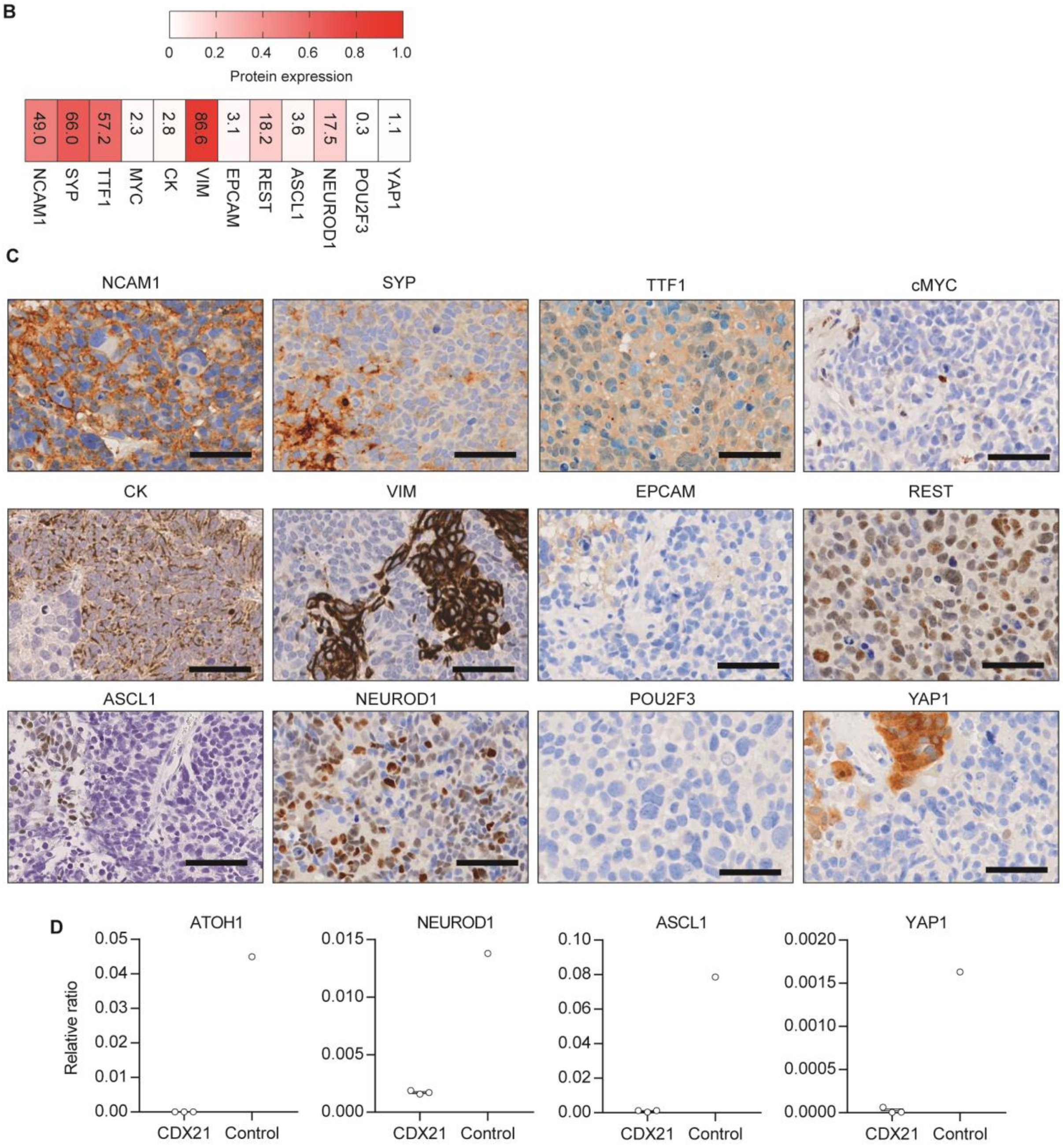

**Supplementary Figure 1, Related to Figure 1. Clinical Features of CDX21 donor patient. (A)** Clinical Features of CDX21 donor patient. **(B)** Heatmap of SCLC diagnostic biomarker expression in CDX21. For each Immunohistochemistry (IHC) assay a whole-tumor section from 3 mice (2 for VIM and EPCAM) was analyzed and the mean expression value was generated in HALO. Mean average % SCLC biomarker expression shown within heatmap. **(C)** Representative IHC images (brown stain). Scale bars, 50 µm. **(D)** RT Real-time quantitative PCR analysis of *ATOH1*, *ASCL1*, *NEUROD1* and *YAP1* in CDX21 tumors compared to a positive control. Relative ratio to *ACTB* control, mean values are shown (black lines) where each circle represents one independent analysis, error bars are plus/minus Standard Error of Mean (S.E.M).

**Supplementary Figure 2, Related to Figure 1. Perfused VM and endothelial vessels are present in SCLC CDX.**

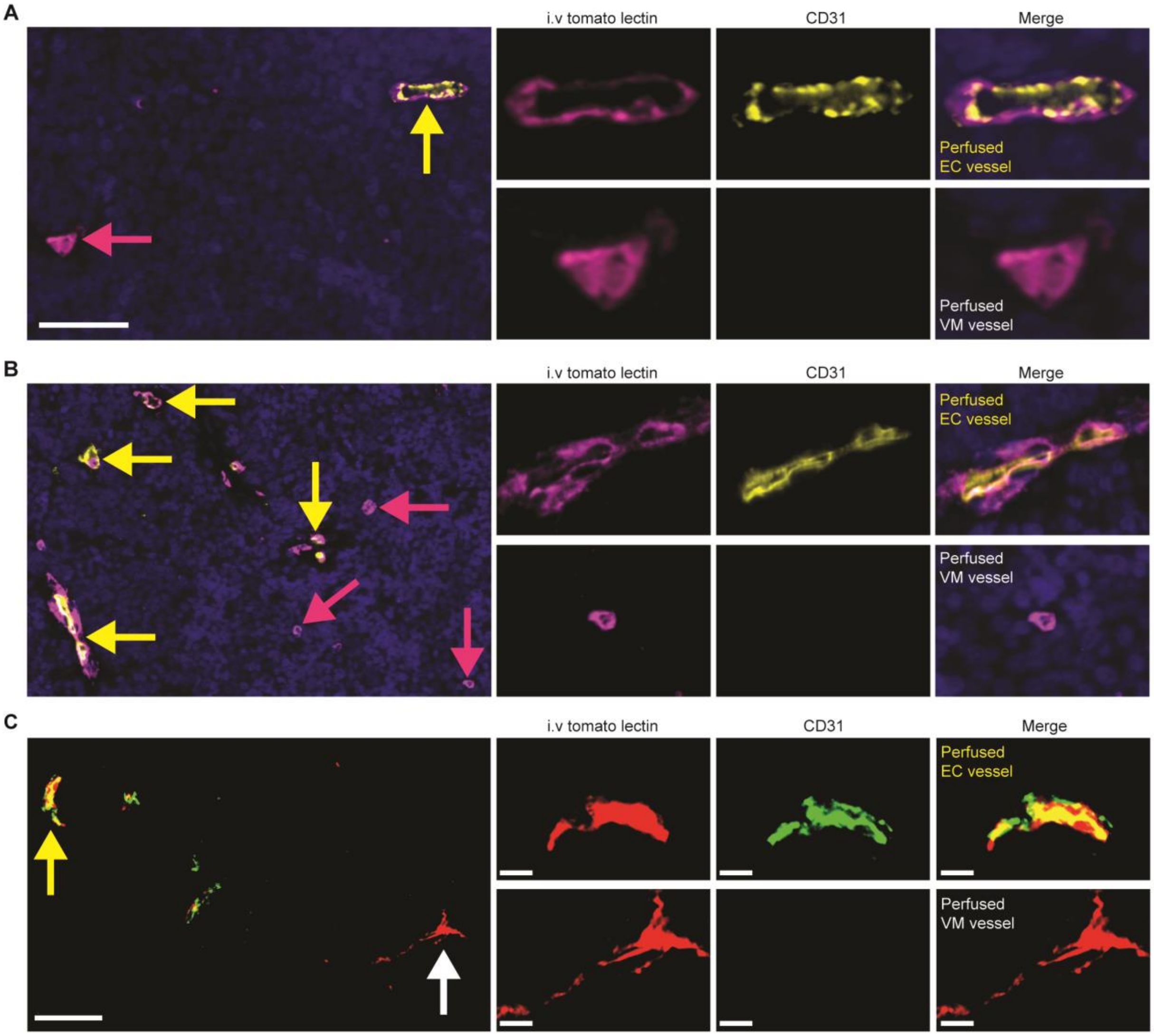

**Supplementary Figure 2, Related to Figure 1. Perfused VM and endothelial vessels are present in SCLC CDX. (A, B)** Representative immunofluorescence image of a perfused endothelial (EC) vessel (CD31$^{+}$/intravenous (i.v) tomato lectin$^{+}$, yellow arrow) and a perfused VM vessel (CD31$^{-}$/i.v tomato lectin$^{+}$, pink arrow) in an FFPE CDX09 tumor with i.v tomato lectin injection. Single channel IF for CD31 (yellow) and i.v tomato lectin (pink) shown with merged multiplex on the right where DAPI is blue. Scale bars 50 µm (left panel) and 10 µm (right panels). **(C)** Representative immunofluorescence image of a perfused endothelial (EC) vessel (CD31$^{+}$/intravenous (i.v) tomato lectin$^{+}$, human mitochondria$^{--}$) and a perfused VM vessel (CD31$^{-}$/i.v tomato lectin$^{+}$, human mitochondria$^{+}$) in a cryosectioned CDX09 tumor with i.v tomato lectin injection. Single channel IF for CD31 (green), i.v tomato lectin (red) and human mitochondria (white) shown with merged multiplex on the right where CD31 and lectin overlay is yellow. Scale bars 10 µm.

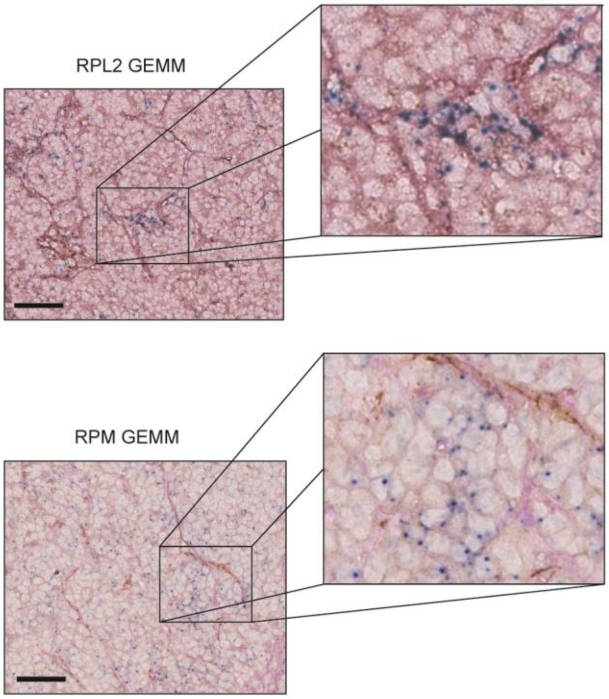


**Supplementary Figure 3, Related to Figure 2. VM co-localizes with REST in SCLC GEMMs.** Multiplex Immunohistochemistry (IHC) and *in situ* hybridization (ISH) assay showing images of VM vessels ($PAS^+/CD31^-$, pink), endothelial vessels ($PAS^+/CD31^+$, brown) and REST ISH (blue). Scale bars, 50 µm. Representative images from three independent tumor replicates in SCLC GEMMs; RBL2 GEMM ($Trp53^{fl/fl}/Rb1^{fl/fl}/Rbl2^{fl/fl}$) [22] and RPM GEMM ($Trp53^{fl/fl}/Rb1^{fl/fl}/Myc^{LSL/LSL}$) [23].

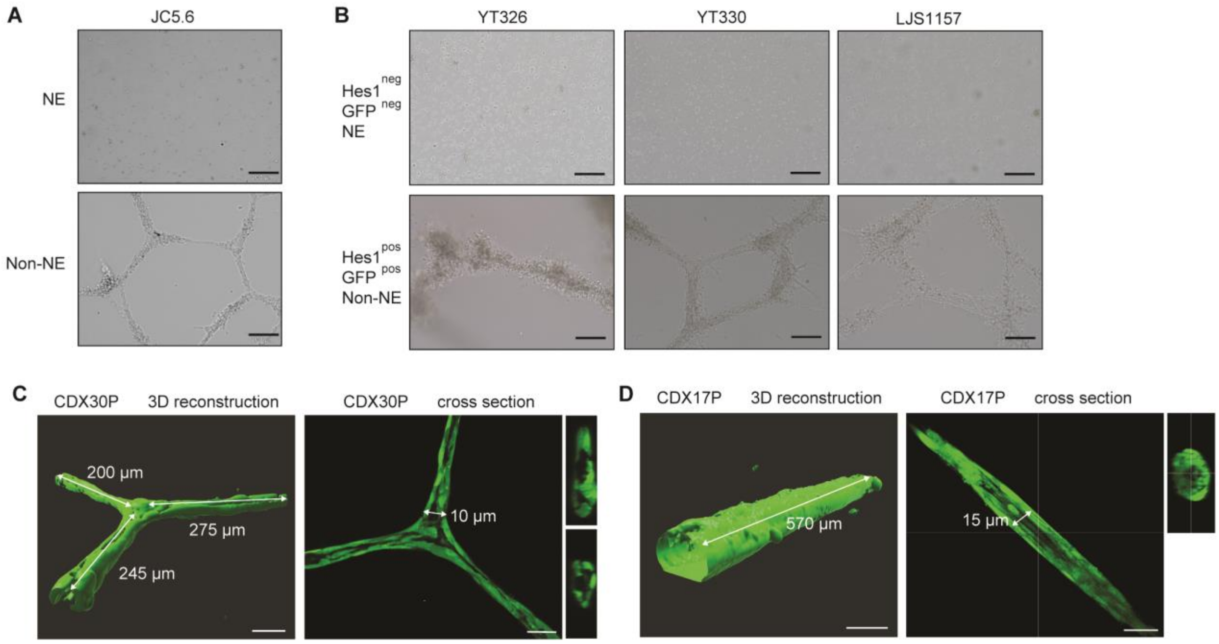


**Supplementary Figure 4, Related to Figure 3. Tubule formation assays in CDX and GEMM tumors *ex vivo.*** **(A)** Representative bright field images of tubule forming assays in RPM GEMM ($Trp53^{fl/fl}$/$Rb1^{fl/fl}$/$Myc^{LSL/LSL}$)[23] NE and non-NE cells *ex vivo*. *n* = 3 independent replicate tumors were analyzed. Scale bars 200 µm. **(B)** Representative bright field images of tubule forming assays in RBL2 GEMM ($Trp53^{fl-/fl}$/$Rb1^{fl/fl}$/$Rbl2^{fl/fl}$)[22] NE ($HES1^{neg}$/$GFP^{neg}$) and non-NE cells ($HES1^{pos}$/$GFP^{pos}$) previously separated by flow cytometry based on *Hes1*-GFP reporter expression and cultured *ex vivo*. *n* = 3 independent replicate cell lines (YT326, YT330 and LJS1157) were analyzed. Scale bars 200 µm. **(C, D)** CDX30P non-NE cells (C) and CDX17P non-NE cells (D) labelled with Cell Tracker Green form hollow tubules when grown on Matrigel. Representative confocal microscopy images followed by Z-stack software reconstruction (Imaris) after 72 hours on Matrigel. Tubule length and diameter dimensions are shown, scale bars 50 µm. CDX30P cross section is the same image as in Figure 3F.

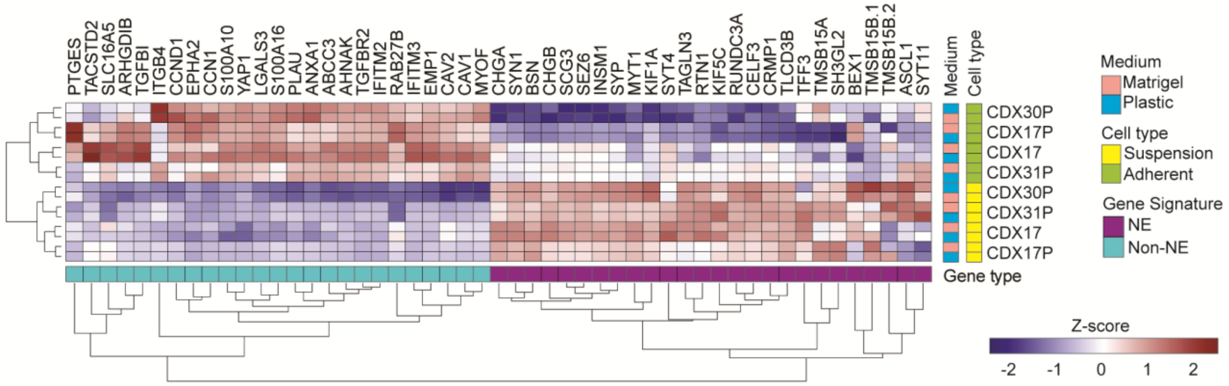


**Supplementary Figure 5, Related to Figure 4. Correlation of NE and non-NE gene and SCLC subtype transcription factor transcript expression in CDX NE and non-NE cells.** 50 gene panel comprising of NE and non-NE genes[10] was mapped to CDX NE and non-NE cell RNAseq data.

**Supplementary Figure 6, Related to Figure 6. RBL2 GEMM ECM adherence assays and CDX PAS/PSR collagen and glycoprotein staining.**

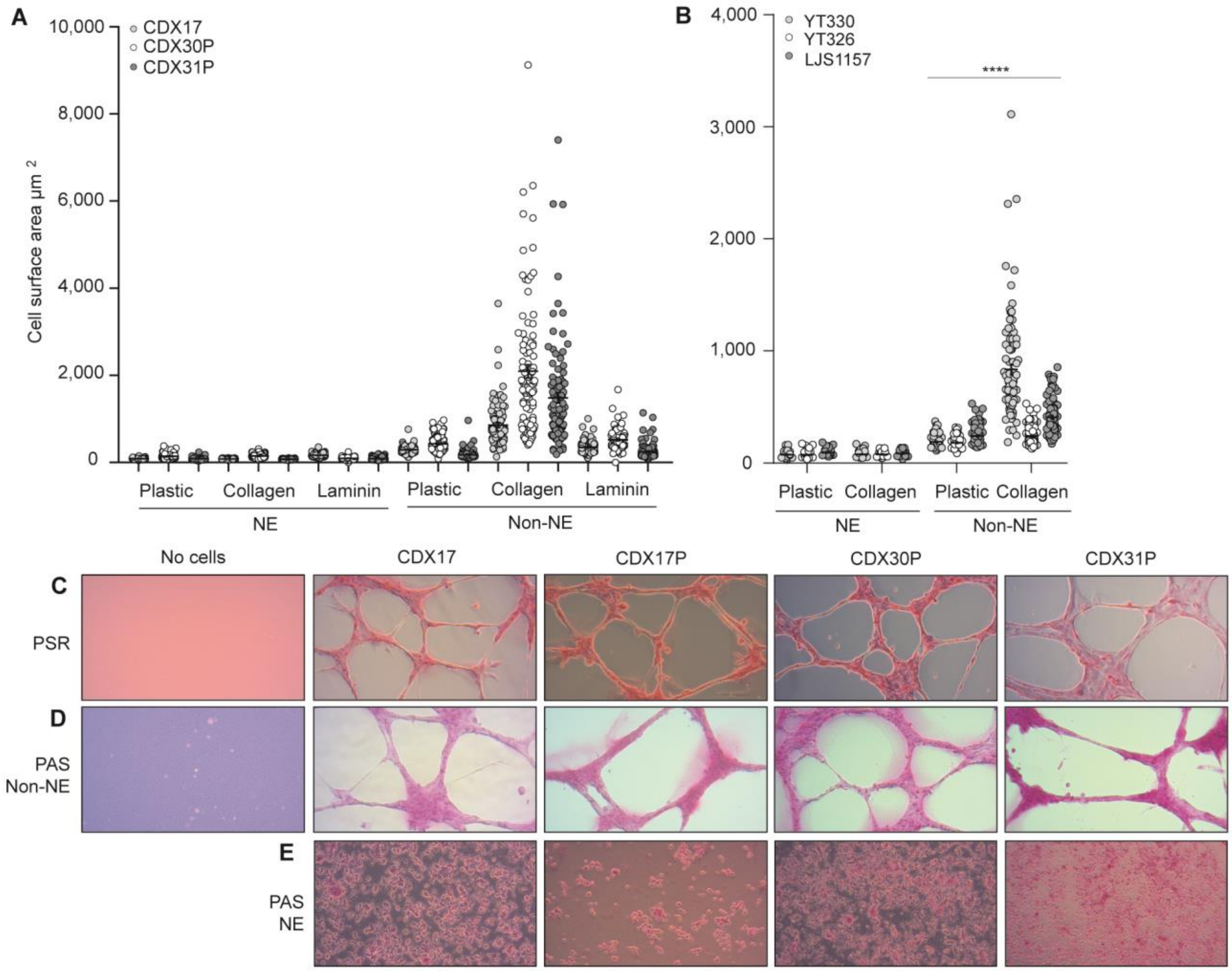

**Supplementary Figure 6, Related to Figure 6. RBL2 GEMM ECM adherence assays and CDX PAS/PSR collagen and glycoprotein staining. (A)** Cell surface area (CSA) of CDX (light gray circles, CDX17, open circles, CDX30P, gray circles, CDX31P) NE and non-NE cells on plastic, collagen, or laminin. Random field of view images were taken and the CSA of 200 cells per CDX tumor was quantified in ImageJ per CDX, represented by each colored circle. Data are mean ± S.E.M. **(B)** Quantification of CSA of RBL2 GEMM ($Trp53^{fl-/fl}$/$Rb1^{fl/fl}$/$Rbl2^{fl/fl}$)[14] NE ($HES1^{neg}$/$GFP^{neg}$) and non-NE cells ($HES1^{pos}$/$GFP^{pos}$) previously separated by flow cytometry based on *Hes1*-GFP reporter expression and cultured *ex vivo* (YT330, light gray circles, YT326, open circles, LJS1157, gray circles) on plastic or collagen. For each assay, random field of view images were taken and the CSA of 200 cells per tumor was quantified in ImageJ. Data are mean ± S.E.M. ****$p < 0.0001$ two-tailed unpaired student's t-test. **(C)** Picrosirius Red (PSR) IHC to detect collagens (red stain) in CDX17, CDX17P, CDX30P and CDX31P non-NE cells (plus acellular control) in *ex vivo* tubule formation assay. **(D)** PAS IHC to detect glycoproteins (pink stain) in CDX17, CDX17P, CDX30P and CDX31P non-NE cells (plus acellular control) in ex vivo tubule formation assay. Representative images shown, *n* = 2-3 samples analyzed **(E)** PAS IHC to detect glycoproteins (pink stain) in CDX17, CDX17P, CDX30P and CDX31P NE cells VM in *ex vivo* tubule formation assay. Representative images shown, *n* = 2-3 samples analyzed per CDX.

**Supplementary Table 1, Related to Figure 1 and Supplementary Figure 1. SCLC subtype classification (ASCL1, NEUROD1, POU2F3 and ATOH1) [8] and YAP1 RNA and/or protein expression [27] of the CDX models used in this study.**

| CDX | Subtype (Simpson et al., 2020) | YAP1 (Pearsall et al., 2020) |
|---|---|---|
| CDX01 | ASCL1 | RNA |
| CDX02 | ASCL1 | |
| CDX03 | ASCL1 | |
| CDX03P | ASCL1 | |
| CDX08 | NEUROD1 | |
| CDX08P | NEUROD1 | |
| CDX09 | ASCL1 | |
| CDX10 | ASCL1 | |
| CDX12 | ASCL1 | Protein |
| CDX13 | POU2F3 | |
| CDX14P | ASCL1 | |
| CDX15P | ASCL1 | RNA and protein |
| CDX15PP | ASCL1 | RNA and protein |
| CDX17 | ATOH1 | |
| CDX17P | ATOH1 | |
| CDX18 | ASCL1 | RNA |
| CDX18P | ASCL1 | RNA |
| CDX20 | ASCL1 | |
| CDX20P | ASCL1 | |
| CDX21* | NEUROD1 | Protein |
| CDX22P | ASCL1 | RNA and protein |
| CDX25 | ATOH1 | RNA |
| CDX29 | NEUROD1 | |
| CDX30P | ATOH1 | RNA and protein |
| CDX31P | ASCL1 | RNA and protein |

P refers to a model made from a blood sample taken during post therapy disease progression. PP refers to model derived from a second post chemotherapy longitudinal blood sample. * denotes CDX21, a newly characterized model used and phenotyped in this study (Figure S1).

**Supplementary Table 2, Related to Figure 4. NOTCH pathway and MYC family member expression in CDX NE versus non-NE cells.**

| Gene | Fold change NE vs non-NE | Adjusted p-value |
|---|---|---|
| **Upregulated in non-NE** | | |
| NOTCH2 | 5.4 | 7.4E-26 |
| NOTCH3 | 12.8 | 3.1E-36 |
| HES1 | 2.2 | 2.5E-06 |
| MYC | 9.3 | 4.5E-19 |
| **Upregulated in NE** | | |
| DLL1 | 22.0 | 6.2 E-18 |
| DLL3 | 4.2 | 1.6E-12 |
| DLL4 | 13.0 | 9.1E-14 |
| MYCL | 14.2 | 2.1E-07 |

Notch pathway receptors (*NOTCH2* and *NOTCH3*), NOTCH pathway ligands (*DLL1, DLL3* and *DLL4*), NOTCH effector (*HES1*) and MYC family member (*MYC, MYCL)* transcript fold change and adjusted p-values in CDX NE versus non-NE cells.

**Supplementary Table 3.xml, Related to Figure 4. Endothelial specific, blood vessel development, angiogenesis and coagulation that are significantly up-regulated in CDX non-NE cells**

**Supplementary Table 4. Antibodies used for immunohistochemistry.** Ready to use (RTU)

| Antibody | Company | Dilution | Antigen Retrieval | Incubation |
|---|---|---|---|---|
| REST | ThermoFisher /MA5-24606 | 1:150 | pH6 20’ | 60’ |
| Synaptophysin | Leica Biosystems/pA0299 | RTU | pH9 20’ | 20’ |
| CD31 | Abcam/ab124432 | 1:400 | pH 6 20’ | 20’ |
| CD31 (IF) | Cell Signaling/77699 | 1:200 | pH 6’20’ | 30’ |
| ASCL1 | BD Pharminigen/556604 | 1:250 | pH6 20’ | 20’ |
| NEUROD1 | Abcam/ab213725 | 1:100 | pH6 10’ | 20’ |
| YAP1 | Abcam/ab52771 | 1:100 | pH6 20’ | 20’ |
| POU2F3 | Sigma-Aldrich/HPA0196652 | 1:250 | pH6 20’ | 20’ |
| NCAM1 | Leica Biosystems/CD56-504-L-CE | 1:100 | pH6 20’ | 20’ |
| TTF1 | DAKO/M3575 | 1:200 | pH9 40’ | 16’ |
| cMYC | Abcam/ab32072 | 1:75 | pH6 20’ | 30’ |
| Cytokeratin | Dako/M3515 | 1:100 | pH6 20’ | 20’ |
| Vimentin | Ventana/790 2917 | RTU | pH9 32’ | 16’ |
| VCAM1 | Abcam/ ab134047 | 1:1000 | pH6 20’ | 30’ |
| EpCAM | Cell Signaling/2929 | 1:100 | pH6 30’ | 20’ |

**Supplementary Table 5. Antibodies used for Immunoblotting**

| Antibody | Company | Clone | Dilution |
|---|---|---|---|
| ASCL1 | BD Pharminigen/556604 | 24B72D11.1 | 1:500 |
| NEUROD1 | Abcam/ab213725 | EPR20766 | 1:500 |
| Synaptophysin (SYP) | Abcam/ab32127 | YE269 | 1:20000 |
| REST | LifeSpan Biosciences/LS-C668231 | N/A | 1:500 |
| YAP1 | Abcam/ab52771 | EP1674Y | 1:1000 |
| NOTCH1 | Bethyl laboratories/A301-895A | N/A | 1:500 |
| NOTCH2 | Bethyl laboratories/A302-083A | N/A | 1:500 |
| cMYC | Abcam/ab32072 | Y69 | 1:500 |
| Vimentin | Cell Signaling Technologies/CST5741 | D21H3 | 1:500 |
| HIF-1a | Abcam/ab51608 | EP1215Y | 1:500 |
| CA9 | Novus Biologicals/NB100-417 | N/A | 1:500 |
| GLUT1 | Abcam/ab115730 | EPR3915 | 1:10000 |
| COL1A1 | Abcam/ab34710 | N/A | 1:1000 |
| ITGA11 | R&D Systems/AF4235 | N/A | 1:500 |
| AXL | Cell Signalling Technologies/CST8661 | C89E7 | 1:500 |
| VCAM1 | R&D Systems/BBA5 | BBIG-V1 | 1:500 |
| PCOLCE | R&D Systems/MAB2627 | 261730 | 1:500 |
| CD44 | Abcam/ab157107 | N/A | 1:1000 |
| Tubulin | Cell Signaling Technologies/CST2144 | N/A | 1:1000 |
| GAPDH | Cell Signaling Technologies/CST2118 | 14C10 | 1:1000 |
| FAK | BD Biosciences, 610088 | 77/FAK | 1:150 |
| Phospho-FAK (Tyr397) | ThermoFisher Scientific, 44624G | N/A | 1:500 |
